# Selection of the Optimal Long-Acting Injectable Formulation of Ivermectin for use in humans to Target Malaria Vectors in Western Africa: Evaluation of Pharmacokinetics and Mosquitocidal Efficacy in Cattle under Laboratory Conditions

**DOI:** 10.1101/2025.07.23.665952

**Authors:** Lamidi Zela, Sié Hermann Pooda, Angélique Porciani, Samuel Beneteau, André Barembaye Sagna, Sophie Le Lamer-Déchamps, Nicolas Moiroux, Cheick Oumar Wendpagnandé Ouédraogo, Anyirékun Fabrice Somé, Cédric Pennetier, Christophe Roberge, Adrien Marie Gaston Belem, Koumbobr Roch Dabiré, Guiguigbaza-Kossigan Dayo, Karine Mouline

**Author notes:** ASTRE, CIRAD-INRAE, Univ Montpellier, Centre de Recherche et de Veille sur les Maladies Vectorielles dans la Caraïbe (CRVC), Petit Bourg, Guadeloupe, France.

## Abstract

**Background:** Ivermectin, a semisynthetic endectocide, is widely used against parasitic nematodes in humans and animals. Its lethality to *Anopheles* mosquitoes after feeding on treated hosts represents a promising malaria control strategy, particularly against outdoor transmission. However, standard oral formulations for use in humans produce short-lived mosquitocidal blood concentrations, limiting epidemiological impact. To meet WHO Preferred Product Characteristics (PPC) for endectocides against malaria vectors (Hazard Ratios >4 for at least one month), three long-acting injectable ivermectin formulations (LAIFs) based on BEPO® depot technology were developed and compared in cattle to identify the most suitable candidate for future human use.

**Methods:** A cattle–*Anopheles* model was used under laboratory conditions in Bobo-Dioulasso, Burkina Faso. Three LAIF candidates (mdc-STM-001, mdc-STM-002, mdc-STM-003) were injected to calves (n=5 per formulation) at 0.6 mg/kg, with untreated calves as controls (n=5). Plasma ivermectin concentrations were measured over 130 days and analyzed using non-compartmental pharmacokinetics. Direct skin feeding assays were conducted at 15 timepoints (days 2–126 post-injection) using insecticide-susceptible (KIS) and wild-derived resistant (VK5) *Anopheles* colonies. Efficacy was assessed through 10-day cumulative mortalities, hazard ratios, 50% lethal concentrations (LC50), and duration of exposure above the 10-day LC50, accounting for the extrinsic incubation period of *Plasmodium falciparum*.

**Results:** All formulations were well tolerated. Mdc-STM-001 showed the most favorable pharmacokinetic profile, with a controlled peak concentration (Cmax = 34.5 ± 12.7 ng/mL) and the lowest inter-individual variability (12%). Ten-day hazard ratios exceeded 4 and cumulative mortalities were >50% for at least 60 days in both strains. Median mosquito lifespan remained below 10 days for at least 90 days post-injection. The 10-day LC50 for resistant mosquitoes (3.66 [2.69–4.97] ng/mL) was maintained for ≥126 days.

**Conclusion:** The Mdc-STM-001 was identified as the optimal candidate. A single injection induced sustained mosquitocidal efficacy for at least two months, achieving HR >4 against both susceptible and resistant *Anopheles* populations and meeting WHO PPC for malaria endectocides. Although extrapolation from cattle to humans requires caution, the favorable pharmacokinetic profile and robust entomological outcomes support progression to Phase 1 clinical trials. Ivermectin’s established safety record further strengthens the rationale for clinical development.

## Background

Malaria remains a leading global health concern with an estimated 263 million clinical cases and 597,000 deaths in 2023, over 95% of which occurred in Sub-Saharan Africa [1]. Significant gains in the fight against the disease have been made between 2000 and 2015 [2] mainly due to indoor residual spraying of houses and universal coverage with pyrethroid insecticide treated nets, two core vector control interventions recommended by the World Health Organization (WHO) [3]. Progresses have stalled since then[4], notably coinciding with widespread, growing, resistance in *Anopheles* vectors populations, due to the huge selection pressure exerted by continuous insecticide-based interventions and massive pesticide use in agricultural practices [5]. Resistance refers to both (i) physiological resistance, involving reduced susceptibility to the insecticidal molecule through target-site mutations or metabolic detoxification processes [6], and (ii) behavioral (qualitative) resistance, as defined by Carrasco et al. [7], whereby mosquitoes avoid insecticides exposure spatially (by biting outdoors), temporally (by biting early in evening or morning when humans are outside their nets or housing), or trophically (by feeding on non-human hosts). New generations of Long Lasting Impregnated Nets (LLINs) incorporating dual active ingredients or synergists [8] [9] have been recently recommended by the WHO to circumvent pyrethroid resistance [10]. However, these tools still primarily target nocturnal, indoor-biting *Anopheles* mosquitoes, and are logically bypassed by behaviorally resistant vectors [11,12]. The resurgence in malaria cases in recent years [1] highlights the urgent need for novel and complementary strategies to reinforce existing interventions and reverse this concerning trend. In particular, tools targeting the diverse biting behaviors of *Anopheles* vectors, whether innate or induced by interventions, are essential for addressing this challenge [7,13].

Mass drug administration (MDA) to humans of the ivermectin endectocide is a proposed approach to overcome limits of current malaria vector control methods [14,15]. Acting systemically, the molecule targets internal parasites as well as external blood-feeding arthropods, including *Anopheles* malaria mosquitoes. It thus primarily reduces *Anopheles* survival probability, the most critical determinant of their vectorial capacity [16]. Because the insecticide is delivered through the host’s bloodstream, it functions regardless spatial or temporal mosquitoes blood-feeding adaptations. Ivermectin is the leading endectocide candidate for vector control notably because of its strong, well established, safety profile from nearly 4 decades of extensive clinical use in MDA campaigns against lymphatic filariasis and onchocerciasis [17]. This safety reflects its selective action on glutamate-gated chloride channels unique to invertebrates [18]. Such mechanism of action is distinct from those of the different insecticide classes used in public health against malaria vectors, so ivermectin is a potent vector control tool that could mitigate current physiological resistance issues. All *Anopheles* species and strains tested to date have been shown to be susceptible to ivermectin, although the degree of susceptibility varies among species [19]. Moreoever sub-lethal effects are also interesting to transmission control as they negatively impact vectors fecundity and fertility [20,21], locomotor and biting behaviors [22,23], or *Plasmodium* parasite development [24,25]. Mass drug administration of ivermectin to humans may therefore effectively reduce residual malaria transmission.

Various oral drug dose regimens have been used during Randomized Controlled Trials (RCTs) testing ivermectin MDAs against malaria. Standard oral ivermectin dosing for parasitic disease treatment in humans (150–200 µg/kg) achieves lethal plasma levels for *Anopheles* mosquitoes for fewer than seven days [26]. In field studies from Sub-Saharan Africa, this regimen administered every three weeks during the rainy season to populations in southern Burkina Faso resulted in a significant 20% reduction in malaria incidence among children under five [27]. Increased dosing regimens (1×400 and 3×300 µg/kg) prolong ivermectin persistence in the host bloodstream and mosquitocidal efficacy up to 14-21 with good safety [28,29]. Modeling studies have predicted that 80% coverage of the target population using such regimen has the potential to reduce malaria incidence by 30 to 70% depending on the transmission settings [30].

In the fields, the BOHEMIA RCT in Mozambique and Kenya applied the 1X 400 µg/kg regimen monthly for three consecutive months across the rainy season [31]. In Kenya, this led to reduced malaria incidence in children aged 5-15 years by 26% compared to the control arm [32]. However, in Mozambique, the same regimen failed to produce significant reduction [33]. Two other RCTs testing the 3 × 300 µg/kg dose administered monthly in combination with either dihydroartemisinin–piperaquine (DHAP) in the Galápagos Islands (MATAMAL [34]) or seasonal malaria chemoprevention (SMC) in Burkina Faso (RIMDAMAL II [35]) also showed no significant epidemiological effect. Possible explanations include biological factors related to vector populations such as poor translation of mosquitocidal effects from laboratory studies to the field or overlooked aspects of wild mosquito bionomics in the trial [36]. Pharmacokinetic data from the RIMDAMAL II trial revealed also a shorter-than-expected residence time of lethal plasma ivermectin, as drug levels dropped below detection in nearly 50% of participants by day 14 [35]. In Mozambique, logistical challenges increased the administration rounds duration so that population coverage never reached the threshold expected for efficacy. All trials have however demonstrated safety of increased and repeated dosing of oral ivermectin.

Both pharmacokinetic limitations and operational challenges encountered in the fields are intrinsic to oral dosing and highlight the need for long-acting ivermectin formulations that can sustain mosquitocidal effects while allowing logistical flexibility. To date, only a single human-targeted long-acting formulation has been developed a gastric-resident oral device with approximately 14-day efficacy [37].

In this context, using the BEPO® injectable technology [38], we developed three ivermectin LAIFs for human use in malaria endemic regions. These formulations were designed according to the efficacy targets set by the WHO recommendations for preferred product characteristics of endectocides in malaria vector control [39], which specify a hazard ratio (HR) greater than 4 for mosquitoes feeding on treated hosts for at least one month post-treatment.

We used a cattle-mosquito colony model [21,40], exposing both insecticide-susceptible and pyrethroid-resistant *Anopheles* strains through direct skin feeding assays, thereby assessing efficacy despite potential cross-resistance or tolerance mechanisms. Our main objective was to compare pharmacokinetic (PK), pharmacodynamic (PD), and mosquitocidal efficacy outcomes among the three candidate formulations to identify the optimal one for subsequent clinical evaluation in human.

## Methods

### Study site

All experiments were conducted in Bobo-Dioulasso, south-western Burkina Faso, through a collaboration between the Institut de Recherche en Sciences de la Santé (IRSS) and the Centre International de Recherche-Développement sur l’Élevage en zones Subhumides (CIRDES). Both institutes have controlled insectary facilities, and CIRDES also maintains stables for cattle husbandry.

### Mosquito colonies

Two *Anopheles* mosquito colonies were used in this study:

(i) ***Anopheles (An.) gambiae* Kisumu (KIS)**: fully susceptible to all classes of insecticides and maintained in the IRSS insectaries. The colony is routinely monitored for contamination by other species and screened for insecticide resistance using molecular techniques [41,42]. For the experiments, mosquitoes were transported in meshed cages covered with moist cloth to the Centre International de Recherche-Développement sur l’Élevage en zone Sub-humide (CIRDES). Transport duration was approximately 10 minutes.
(ii) ***An. coluzzii* VK5 (VK5)**: This colony was established at CIRDES in November 2020, prior to the start of the experiments in January 2021. A total of 150 blood-fed females were collected from the village of VK5 in the Kou Valley (∼50 km northwest of Bobo-Dioulasso), an irrigated rice growing area where different classes of pesticides are used for agriculture [43]. The use of pyrethroid impregnated bed nets is high due intense mosquito nuisance. The mosquito’s bio-ecology is well characterized, with *An. coluzzii* representing more than 95% of *Anopheles* species, and permanently found breeding in the irrigated fields. *An. coluzzii* populations are there characterized by a high prevalence of resistance to all the insecticides classes, justifying the choice of this cite to found the insecticide resistant colony. Collected females were allowed to individually oviposit. Species identification was performed post-oviposition by PCR using established protocols [41] and only progeny from confirmed *An. coluzzii* were retained. As expected, based on the collection site, all females belonged to the target species.

Both colonies were housed in similar environmental conditions and maintenance protocol [40]. The insectaries temperature was set to 26°C ± 1°C, relative humidity to 75% ± 5%, and photoperiod to 12-hour light/dark. The VK5 colony’s pyrethroid resistance status was routinely monitored by sampling approximately 40 females per generation and screening for the knockdown resistance West African allele (KDR-W; L1014F) of the voltage-gated sodium channel gene using quantitative polymerase chain reaction (qPCR) [42]. For each direct skin-feeding assay time point, females were randomly drawn from at least five different cages to minimize potential cage effects.

### Long acting ivermectin formulations

The long-acting ivermectin formulations tested in this study were developed using BEPO® technology [38], a proprietary long-acting injectable technology (Medincell, Jacou, France). Following subcutaneous injection, BEPO® forms an *in situ* bioresorbable polymeric depot via a solvent-exchange mechanism, leading to precipitation of a polymer matrix that enables sustained drug release. Each formulation consisted of three components: i) a biocompatible solvent to ensure injectability, ii) ivermectin, as the active pharmaceutical ingredient, and iii) a mixture of two biocompatible and biodegradable polymers that control drug release kinetics. These polymers undergo progressive hydrolysis into bioabsorbable by-products over time, thanks to a simple hydrolysis mechanism. Three ready-to-use LAIFs solutions were evaluated: mdc-STM-001, mdc-STM-002 and mdc-STM-003; the last one differed in ivermectin loading and each differed in polymer composition and proportions.

All formulations were imported into Burkina Faso with clearance from the Direction Générale des Services Vétérinaires of Burkina Faso (Permit No 2020/199) and were stored at room temperature, protected from light, until administration to cattle. The study protocol was approved by the Institutional Animal Ethics Committee of CIRDES (document reference number: 004-10/2020 CEB-CIRDES).

### Cattle hosts and treatments

Twenty-five male calves (Métis strain: Fulani zebus x Baoulé crossbreed) with a median weight of 117 kg (interquartile range [IQR]: 106-129 kg), were purchased from villages in the Valley du Kou area two months prior to the study to allow for acclimation. Upon arrival at the CIRDES animal facility, calves were treated for trypanosomiasis and gastrointestinal parasites using Berenil 2000® and Benzal®, respectively. Animals were housed outdoors under screened enclosures to prevent exposure to or reinfestation with ectoparasites and trypamosomoses. They were fed with rice straw and 1 kg/day of cottonseed cake, with water and sea salt provided ad libitum.

Calves were randomly assigned to five experimental arms (n = 5 calves per arm) using stratified randomization to balance body weight across groups. Four arms received LAIFs, and one served as the untreated control. The formulations were administered *via* a single subcutaneous injection into the loose skin anterior to the shoulder.

- mdc-STM-001 and mdc-STM-002 LAIFs were each injected at a unique dose of 0.6 mg/kg (referred to as mdc-STM-001-0.6 and mdc-STM-002-0.6, respectively).
- mdc -STM-003 was administered at two doses: 0.6 mg/kg and at 1.5 mg/kg (referred to as mdc-STM-003-0.6 and mdc-STM-003-1.5).

The 1.5 mg/kg dose represents the upper limit of feasible subcutaneous volumes for human application and was included to assess dose-dependent effects on pharmacokinetics and mosquitocidal efficacy. Nominal injection volumes per animal ranged from 0.62 to 2.14 mL (detailed in Table 1). The animals’ identification numbers, their weight upon arrival at CIRDES and on the day of injection, administered formulations, and injected volumes are provided in the Additional file 1, Table S1. Animal health was monitored throughout the study by assessing each day clinical signs, local injection site reactions, body weight, and weight gain.

**Table 1:**
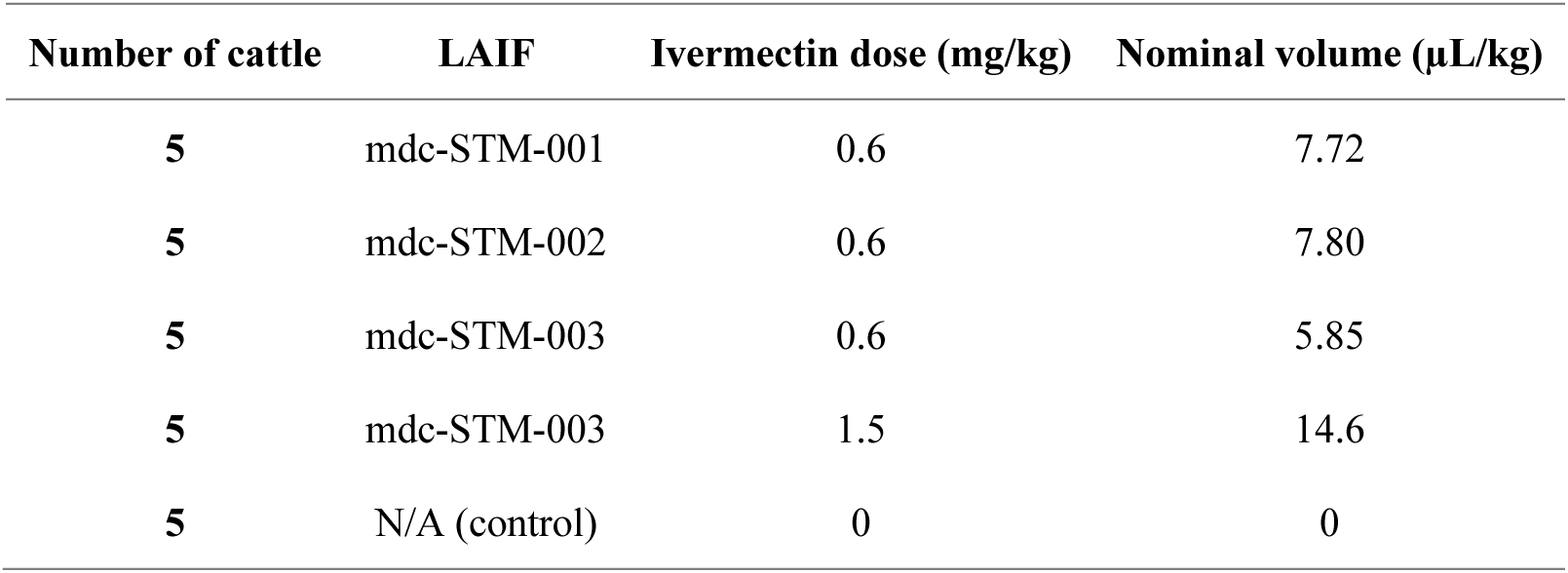
Nominal treatment volumes per formulation and dose to be administered to cattle. N/A: not applicable.

### Mosquito blood-feeding bioassays

To ensure equivalent ivermectin exposure, mosquitoes from both KIS and VK5 colonies were simultaneously exposed to the same cattle hosts during blood-feeding. This design enabled direct comparison of susceptibility across colonies under identical conditions. Direct skin blood-feeding assays were conducted following previously established protocols [40]. Prior to feeding, mosquitoes were deprived of sugar for 12 hours, with access to water-soaked cotton only. At each time point, approximately 100–130 females per colony were introduced into net-covered bowls, and held against the shaved flank of the calf for 20 minutes using a rubber band. Separate bowls were used for each colony. Fully engorged mosquitoes were distributed into cups (10 females per cup), with a maximum of 120 cups per colony per time point. All cups were maintained under controlled conditions in the CIRDES insectary. Mosquito mortality was recorded daily between 10:00 and 11:00 a.m. for 30 days post-blood meal. To mitigate positional bias, cups were randomly repositioned on shelves after each observation.

A total of 16 blood-feeding assays were conducted: once pre-treatment (baseline) and at 2, 7, 17, 21, 28, 42, 56, 70, 84, 91, 98, 105, 112, 119, and 126 Days After Injection (DAI). Due to limited mosquito availability, KIS mosquitoes were not tested at 7 and 91 DAI. Surplus blood-fed mosquitoes were stored at –20°C for downstream analyses.

### Ivermectin bioanalysis and pharmacokinetics

Jugular venous blood samples were collected from treated calves at each mosquito exposure time point and at additional time-points (30 samples per animal) to characterize ivermectin pharmacokinetics (PK). Sampling included pre-dose, 2h, 5h and 12h post-injection, then daily sampling until DAI 4, every 2-4 days until DAI 21, and weekly until DAI 132-133. Blood samples were also collected from control animals at pre-dose and at DAI 1, 14, 28, 56, 84 and 111. Blood samples were centrifuged at +4°C (2000 rpm for 10 min) to isolate plasma, which was stored at -80°C until shipment for bioanalytical analysis. Ivermectin plasma concentrations were quantified using a liquid-liquid extraction with qualified LC-MS/MS methods. Two calibration ranges were used (0.2-20 ng/mL low range; 5-500 ng/mL high range), using K_2_-EDTA as anticoagulant and ivermectin-D_2_ as internal standard. The quality controls (0.6, 10, 16 ng/mL low range; 15, 250, 400 ng/mL high range) were within the acceptance criteria (precision ≤ 15.00%, deviation ± 15.00%). Incurred sample reanalysis validated ivermectin stability for at least 680 days at -20°C. Plasma samples collected from cattle were stored at -80°C no longer than 162 days before analysis, validating the ivermectin concentrations measured for PK analysis.

At study completion, control animals were returned to stock, while treated animals were euthanized. Injection sites and depot with surrounding tissues were sampled to quantify the residual ivermectin and copolymers. These analyses were performed using an ultra-performance liquid chromatography (limit of quantification: 0.9 µg/mL) for ivermectin and nuclear magnetic resonance analysis for copolymers. Both methods were developed for exploratory purposes only.

### Outcomes and Analysis

Unless specified otherwise, analysis were performed using the R version 4.4.3 (2025-02-28) [44]. Models fit were assessed using the “performance” [45] and DHARMa [46] packages. All estimates are reported with 95% confidence intervals (CI95%) and/or p-values.

#### Cattle weight gain

Differences in weight gain between treatment groups throughout the study were analyzed using the Kruskal–Wallis rank sum test, followed by pairwise two-sided exact Wilcoxon rank sum tests with 95% confidence intervals. *p*-values were adjusted using the Holm method ([47]).

#### Pharmacokinetics

PK parameters of ivermectin were calculated for each LAIF using a non-compartmental analysis (Phoenix® WinNonlin® version 8.1; Certara USA, Inc., Princeton, NJ). These included the rate of systemic exposure (Cmax), the time to reach Cmax (Tmax), the extent of systemic exposure over the study duration (aera under the curve, AUClast), over 28 DAI (AUC0-28d) and over 91 DAI (AUC0-91d), concentrations at 28 DAI (C28d), 56 days (C56d), 91 DAI (C91d) and at study end (Clast), and the apparent terminal phase half-life (T1/2) when calculable. The systemic exposure parameters were normalized to the actual administered dose, calculated for each cattle based on the weight of each syringe before and after injection. The inter-individual variability in plasma concentrations and PK parameters was expressed as the coefficient of variation (CV %) for each experimental group.

#### Mosquito related outcomes

##### Mosquito blood feeding rate

For each colony, we compared the mosquitoes’ blood-feeding rates across treatment arms using a generalized linear mixed model (GLMM, [48]) with a binomial error distribution. The outcome variable was the number of blood-fed mosquitoes, with the total number of mosquitoes included as an offset. The model included an interaction term between treatment and DAI, as well as a random intercept for cattle. The model was fitted using restricted maximum likelihood.

##### Times post-feeding

Efficacy outcomes are provided for 4- 10 and 30-days post mosquito blood feeding. The 4-day period reflect the acute toxic effect, which typically manifests within 2-3 days post-ivermectin blood meal. The 10-day post-blood feeding period is used in relation the 10-day minimum extrinsic incubation period (EIP) needed for the parasite to develop in the mosquito and to reach the salivary glands [49], thereby rendering it infectious and active in transmission. The 30-day follow-up captures the residual effects of ivermectin on surviving mosquitoes, which may include both indirect sublethal effects and potential interactions with the mosquitoes’ age. Only data related to the 10-day follow up period are presented and thoroughly discussed in the main text. Outcomes for the 4-day and 30-day follow-up period are given in the Additional file 2, Fig. S3 and S4.

##### Cumulative Mortality

Proportions of dead mosquitoes at 10-days post-feeding were computed for each colony, formulation, and DAI.

##### Survival Analysis and Hazard Ratios

Daily mosquito mortality was recorded for up to 30 days post blood-feeding. For each mosquito colony and DAI, survival data were used to generate Kaplan–Meier survival curves, stratified by treatment group.

Per colony, survival patterns at each time-point were compared to the control group using Cox proportional hazards models (mixed-effects regression model [50]). The models included “treatment group” and “DAI” as fixed effects, with random intercepts for individual calf ID and mosquito cup (nested random effects) to account for variability within treatment groups. Hazard ratios (HR) at different time points post-blood feeding are given to assess the impact of ivermectin on mosquito mortality risks and HR variations over time. Post hoc pairwise comparisons of HRs between time points were adjusted for multiple testing using the Tukey method, implemented via the “emmeans” package [44]

##### Dose–Response Analysis and coverage duration

Dose–response relationships were analyzed using the “drc” package in R [51], modeling 4-day and 10-day mosquito mortality as a function of host ivermectin plasma concentrations. A four-parameter log-logistic model was fitted with the upper limit fixed at 1. Confidence intervals were computed using the tfls method (t from logarithm scale).

## RESULTS

### Cattle follow up

A mild reaction (agitation) was observed in most animals after the injection, regardless of dose or LAIF administered (data not shown). There were no unscheduled cattle deaths, and no clinical signs were observed in any of the treated groups throughout the study. The major local reaction was swelling, which occurred in all animals, as expected with this LAIF technology. Median weight gain throughout the study (i.e. 4 months) was unaffected by the test item treatment (Kruskal-Wallis test, χ²= 8.4892, df = 4, p-value = 0.075, Table 2). For each experimental group, monthly weight measures are given in the Additional file 1, Table S2.

**Table 2:** Median cattle weight gain per treatment group with interquartile range (IQR).

|  | Control | mdc-STM-001-0.6 | mdc-STM-002-0.6 | mdc-STM-003-0.6 | mdc-STM-003-1.5 |
| --- | --- | --- | --- | --- | --- |
| Weight gain (kg)<br>Median<br>(Q1-Q3) | 27.4<br>(25.8- 31.6) | 37.2<br>(35.8-44.2) | 39.6<br>(36.6-41.6) | 35.3<br>(31.2- 36.2) | 30.0<br>(28.6-40.0) |

### Ivermectin pharmacokinetics

A total of 635 blood samples were collected, including 600 samples from LAIF-treated cattle (4 treatment groups, 5 cattle per group, n=30 per cattle) and 35 samples from control cattle (1 control group, 5 cattle, n=7 per cattle). Concentration-time patterns were analyzed using data until DAI = 132 for mdc-STM-001-0.6, mdc-STM-002-0.6 and DAI 133 for mdc-STM-003-0.6 and mdc-STM-003-1.5. Ivermectin plasma concentrations were analyzed according to the treatment (formulation and dose). Individual concentration-time profiles of ivermectin after LAIFs injection are illustrated in Figure 1. The mean concentration-time profiles over the study duration and for the first 7 days only to better visualize the burst (i.e., peak plasma level), are provided in the Additional file 2: Fig. S1.

**Figure 1:**
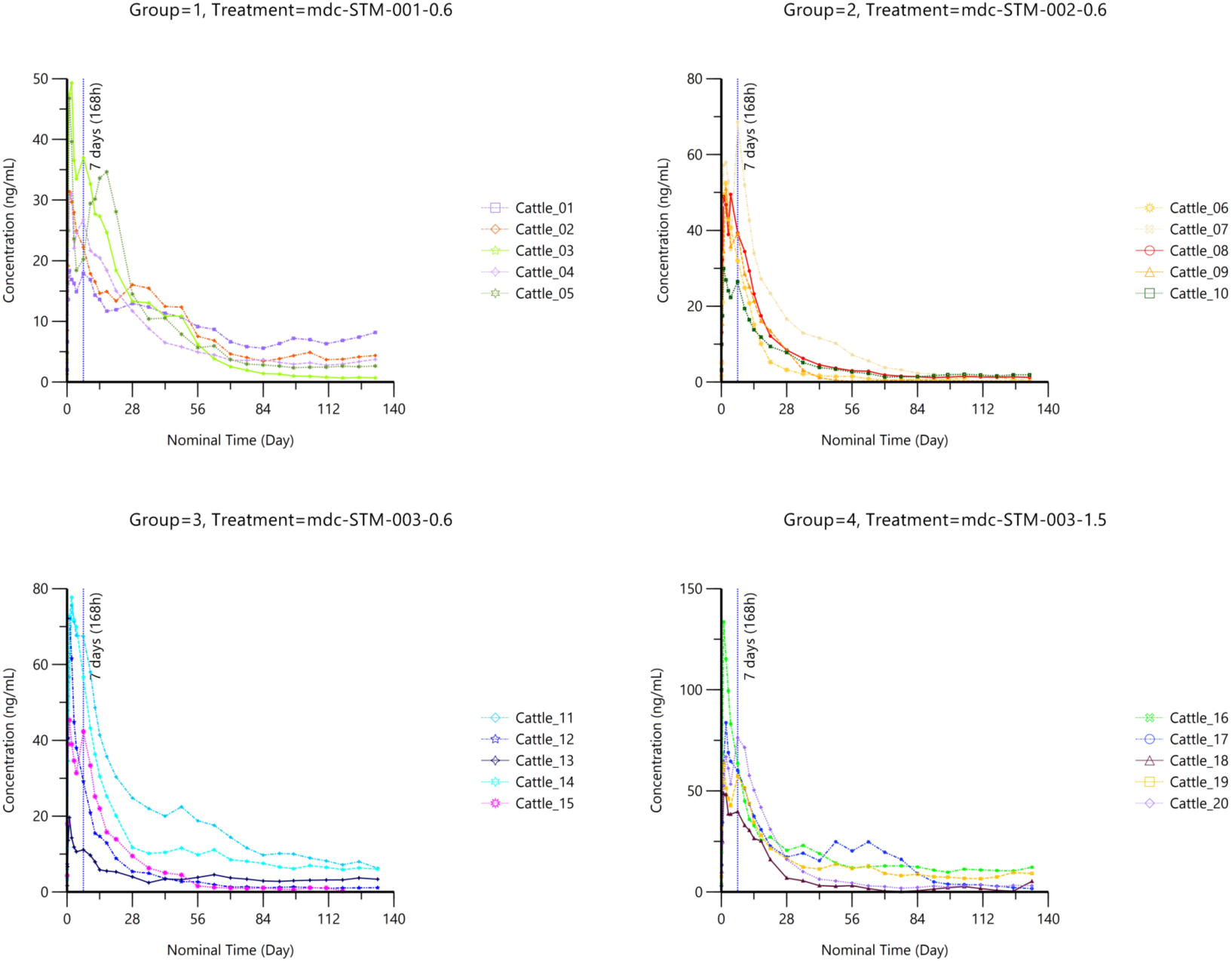
Individual kinetic profiles of ivermectin in cattle plasma after single sub-cutaneous administration of candidate long acting ivermectin formulations (LAIFs) at 0.6 or 1.5 mg/kg

Ivermectin was not detected in any of the samples collected in the control group during the study period nor in any samples collected before ivermectin injection in the treated groups. Following sub-cutaneous injection, ivermectin plasma concentrations were generally measured in all samples collected during the study, with quantifiable levels still present at the final study time-point (*i.e.,* DAI 132 or 133).

After a single sub-cutaneous administration of LAIFs, the plasma levels of ivermectin increased to reach the maximal level (*i.e.,* the burst), known to occur with LAIs [38]) at 24h or 48h post-dose (median Tmax). This was followed by two phases of decrease: a first phase until Day 56 (*i.e.,* 1344h post dose) then a slower phase with a regular decrease of ivermectin plasma levels until the study end. Among the formulations, mdc-STM-001-0.6 showed the lowest rate of decrease during the first phase and the steadiest levels during the second phase. In contrast, the mdc-STM-002-0.6 exhibited the fastest decrease compared to other LAIFs. The inter-animal variability in ivermectin plasma levels (expressed as coefficient of variation CV%) was moderate. The lowest variability was seen with mdc-STM-001-0.6 (mean CV%: 41%, range: 12-70%), followed by mdc-STM-003-1.5 (mean: 54%, range: 22-87). The highest variability was observed for mdc-STM-002-0.6 (mean CV%: 62%, range: 25-106%) and mdc-STM-003-0.6 (mean: 72%, range: 34-98%).

The mean ivermectin PK parameters of each LAIF after injection are reported in Table 3. Additional parameters are given in the Additional file 1: Table S3.

**Table 3:** Non compartmental pharmacokinetic parameters of ivermectin in cattle plasma after single sub-cutaneous administration of the different long acting ivermectin formulations (LAIFs) at a dose of 0.6 (mdc-STM-001, mdc-STM-002, mdc-STM-003) or 1.5 mg/kg (mdc-STM-003). For each parameter, values are expressed as mean ± standard deviation (SD). Tmax: time to reach Cmax. Cmax: maximal plasma concentration. Clast: last quantifiable plasma concentration. AUC(0-28d): area under curve from time 0 to 28 days. AUC(0-28d)/Dose: dose-normalized AUC(0-28d). AUC(0-91d): area under curve from time 0 to 91 days. AUC(0-91d)/Dose: dose-normalized AUC(0-91d). C28d: plasma concentration 28 days post injection. C56d: plasma concentration 56 days post injection. C91d: plasma concentration 91 days post injection.

| Parameters | mdc-STM-001-0.6 | mdc-STM-002-0.6 | mdc-STM-003-0.6 | mdc-STM-003-1.5 |
| --- | --- | --- | --- | --- |
| Tmax <sup>a</sup>(h) | 24 (24; 48) | 48 (24; 168) | 24 (24; 48) | 24 (24; 168) |
| Cmax (ng/mL) | 35.40 ± 12.70 | 50.30 ± 13.70 | 58.10 ± 25.20 | 81.20 ± 32.20 |
| Cmax/Dose<br>(ng/mL/mg/kg) | 58.30 ± 22.00 | 84.30 ± 22.60 | 99.40 ± 42.20 | 56.10 ± 22.40 |
| Clast (ng/mL) | 3.93 ± 2.75 | 1.03 ± 0.70 | 3.43 ± 2.74 | 6.30 ± 4.35 |
| AUC(0-91d)<br>(h.ng/mL) | 25055 ± 2936 | 21121 ± 8866 | 28890 ± 18668 | 41820 ± 13286 |
| AUC(0-91d)/Dose<br>(h.ng/mL/mg/kg) | 40990 ± 5664 | 35479 ± 14839 | 49483 ± 31994 | 28903 ± 9300 |
| C28d (ng/mL) | 13.70 ± 1.60 | 8.90 ± 4.85 | 11.10 ± 8.30 | 15.50 ± 5.10 |
| C56d (ng/mL) | 6.70 ± 1.67 | 2.92 ± 2.63 | 7.34 ± 7.14 | 10.30 ± 6.90 |
| C91d (ng/mL) | 3.48 ± 1.85 | 1.11 ± 0.84 | 4.33 ± 3.99 | 5.55 ± 3.72 |

<sup>a</sup> Median (min; max).

At a dose of 0.6 mg/kg, PK differences emerged across formulations when considering both the entire study duration or the 91-day period (*i.e.,* 3 months post-injection), which is the best targeted release duration. These differences involved performance, quantification levels over time, and inter-animal variability. The peak plasma concentration (Cmax) for mdc-STM-001-0.6 was at least 30% lower compared to the other LAIFs, with mdc-STM-003-0.6 showing the highest Cmax. This suggests that mdc-STM-001-0.6 provided a safer burst compared to the other LAIFs. The inter-animal variability in Cmax was moderate for all formulations, with the lowest variability observed for mdc-STM-002-0.6 (CV% = 27%) followed by mdc-STM-001-0.6 (CV%=36%) and mdc-STM-003-0.6 (CV%=43%). The overall systemic exposure over DAI 91 (AUC_0-91d_) was quite similar for all formulations, with the lowest value for mdc-STM-002-0.6 and the highest value for mdc-STM-003-0.6. Inter-animal variability in AUC was much lower for mdc-STM-001-0.6 (CV% = 12%), compared to mdc-STM-002-0.6 (CV% = 42%) and mdc-STM-003-0.6 (CV = 66%). The mean ivermectin plasma concentrations after each month post sub-cutaneous injection (C28d, C56d, C91d, Clast) decreased slowly until the end of the study for all formulations, remaining close to or above 3.5 ng/mL for mdc-STM-001-0.6 and mdc-STM-003-0.6, and around 1 ng/mL for mdc-STM-002-0.6. There was moderate to high inter-animal variability, ranging from 53% for mdc-STM-001-0.6 to 92% for mdc-STM-003-0.6. Additionally, the ratio of Cmax to the level measured after each month (C28d, C56d, C91d) was up to 2-fold lower for mdc-STM-001-0.6 compared to other LAIFs at the dose of 0.6 mg/kg.

For mdc-STM-003, both tested doses (0.6 and 1.5 mg/kg) exhibited a hypo-proportional increase in Cmax and AUC, approximately 40% less than proportionality. However, these findings should be considered cautiously due to the high inter-animal variability observed, particularly at the low dose (Table 3 and Additional file1: Table S2).

Because the second phase of the PK profile was relatively flat, the apparent terminal phase half-life (T1/2) was accurately estimated for only some animals, with a mean T1/2 ranging from 20 days for mdc-STM-003-1.5 to 33 days for mdc-STM-001-0.6. The remaining amounts of ivermectin depot and copolymers were the lowest in animals treated with the mdc-STM-001-0.6 formulation (Additional file tables S4 and S5).

### Mosquitocidal activity evaluation

#### Knock-down resistance allele (KDR-W) status of the VK5 mosquito lots

Generations F2 to F8 of the VK5 colony were used in this experiment. Mosquito batches exposed at DAI 2, 7, 17, 21, 28, and 42 consisted of 74–80% individuals exhibiting pyrethroid-resistant phenotypes (i.e., homozygous or heterozygous for the KDR-W allele of the voltage-gated sodium channel gene [42]). The remaining batches (exposed at DAI 56, 70, 84, 91, 98, 105, 112, 119 and 126) contained 50–54% resistant mosquitoes. Detailed qPCR results by generation and DAI are given in Supplementary Table S6.

#### Blood feeding rate

High blood-feeding rates were consistently observed across all time points for both KIS and VK5 colonies (Table 4). No statistically significant differences in feeding success were observed between treatment groups or formulations at any time point (data not shown), indicating uniform host attractiveness and feeding opportunity across experimental conditions.

**Table 4:** Blood-feeding rates of KIS and VK5 mosquitoes allowed to feed on cattle from the different treatment groups. Numbers in brackets indicate the number of fed mosquitoes out of the total exposed.

|  | Control | mdc-STM-001-0.6 | mdc-STM-002-0.6 | mdc-STM-003-0.6 | mdc-STM-003-1.5 | Total |
| --- | --- | --- | --- | --- | --- | --- |
| KIS | 0.900<br>(3908/4341) | 0.921<br>(4128/4481) | 0.897<br>(4206/4487) | 0.892<br>(4080/4572) | 0.831<br>(3675/4421) | 0.889<br>(19817/22302) |
| VK5 | 0.966<br>(5047/5224) | 0.945<br>(5005/5295) | 0.935<br>(5091/5447) | 0.961<br>(5160/5371) | 0.968<br>(5174/5343) | 0.955<br>(25477/26680) |

#### Efficacy outcomes

The total number of mosquitoes monitored for survival by colony and treatment group is presented in Table 5. Their detailed distribution by individual cattle, cup, and day after injection (DAI) is provided in Supplementary Tables 6A (KIS) and 6B (VK5). Due to limited availability of KIS mosquitoes, no feeding assays were conducted at DAI 7 and DAI 28. Mosquito husbandry issues affected the same colony at DAI 105 and DAI 119; these data points were excluded from the analysis. Unexpected high mortality rates during the first 4 days post blood feeding (> 0.20) were detected for KIS mosquitoes fed on the control cattle B328 at DAI 42 (mortality rate = 0.437) and B321 at DAI 112 (mortality rate = 0.21). Therefore, these mosquito lots were excluded from efficacy analyses.

**Table 5.** Number of KIS and VK5 mosquitoes followed for survival experiments for each treatment group.

|  | KIS | VK5 |
| --- | --- | --- |
| Control | 1,926 | 3,161 |
| mdc-STM-001-0.6 | 2,122 | 3,206 |
| mdc-STM-002-0.6 | 2,123 | 3,186 |
| mdc-STM-003-0.6 | 2,135 | 3,207 |
| mdc-STM-003-1.5 | 2,093 | 3,182 |
| **Total** | 10,399 | 15,942 |

##### Mortality rates

The mdc-STM-001-0.6 showed the highest efficacy against KIS mosquitoes, comparable to mdc-STM-003-1.5 (Figure 2). From DAI 2 to approximately DAI 100, over 75% of KIS mosquitoes died within 10 days after feeding cattle treated with mdc-STM-001-0.6, and mortality rates remained above 50% beyond this period. For VK5 mosquitoes, mdc-STM-001-0.6 maintained ≥50% mortality between DAI 2 and 80, whereas other formulations at the dose of 0.6 mg/kg were less effective. The high-dose mdc-STM-003-1.5 achieved the longest duration of efficacy for VK5 mosquitoes, with mortality rates >50% from DAI 2 to 110.

**Figure 2.**
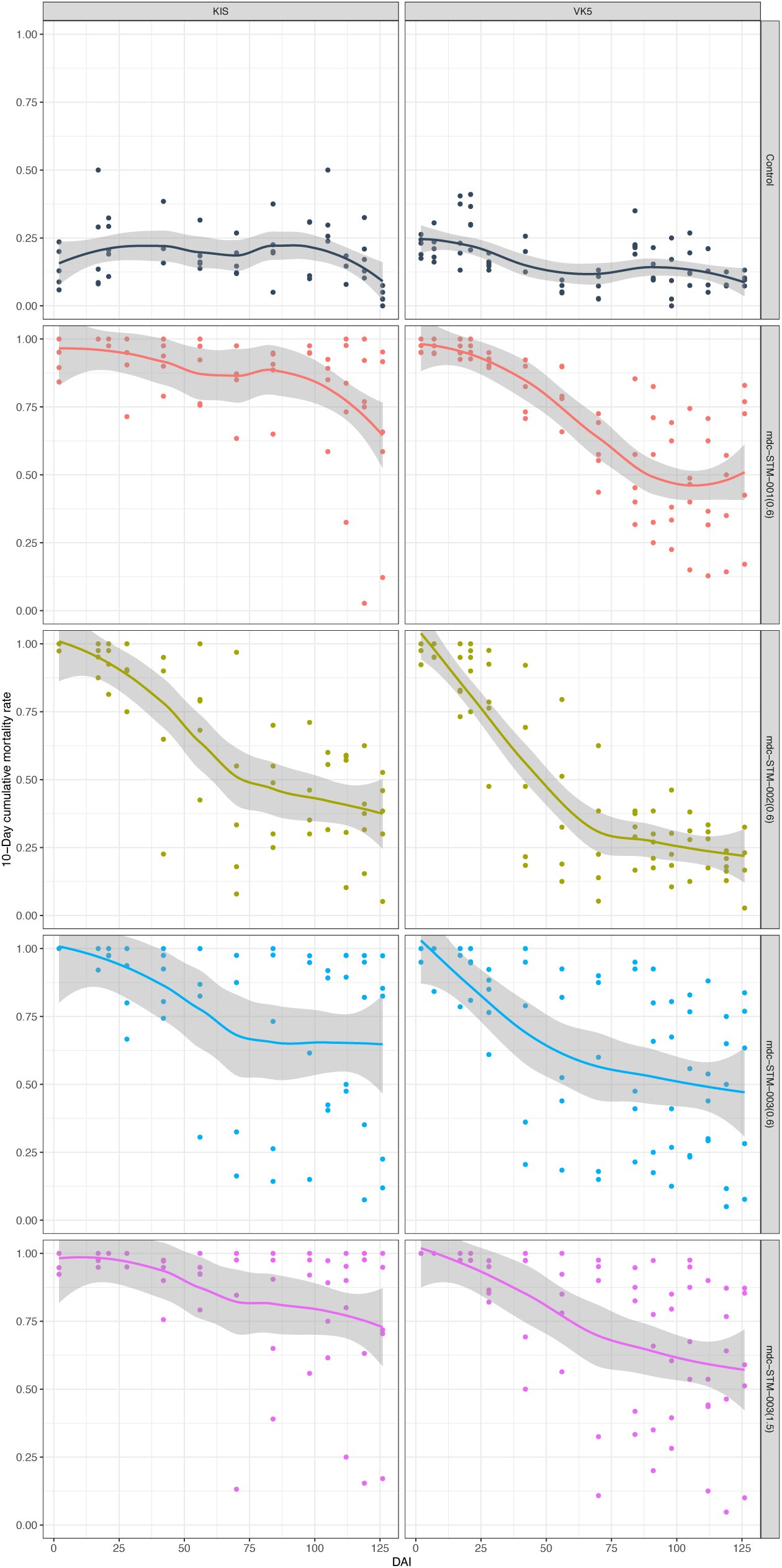
Ten-day cumulative mortality rates of KIS and VK5 mosquitoes fed at different days after injection (DAI) on control or treated cattle. Treated cattle received a single LAIF injection at a dose of 0.6mg/kg for the mdc-STM-001, mdc-STM-002, mdc-STM-003 formulations, and at an additional dose of 1.5 mg/kg for the mdc-STM-003. The smooth line represents mean values estimated by the LOESS (locally estimated scatterplot smoothing) method and grey area shows CI95% [44]. Results for the 4-day cumulative mortalities are shown in the Additional file1: Fig. S3.

Among all formulations, mdc-STM-001-0.6 exhibited the most consistent efficacy profile. Mosquito mortality data for this LAIF showed the lowest inter-animal variability, with fewer deviations from the mean trend across all sampling points.

##### Survival and Hazard Ratios

No significant differences in mosquito survival were observed across treatment groups prior to LAIF administration for either KIS or VK5 colonies (Figure 3, Additional file 1: Table S8, Additional file 2: Fig. S2 and S3), confirming no confounding effects from host variability. Following treatment, all formulations administered at the dose of 0.6 mg/kg (mdc-STM-001, mdc-STM-002, mdc-STM-003) and at 1.5 mg/kg (mdc-STM-003) induced a significant mosquitocidal effect. The magnitude and duration of these effects varied according to mosquito strain and formulation but remained statistically significant through the full follow-up period (DAI 126).

**Figure 3:**
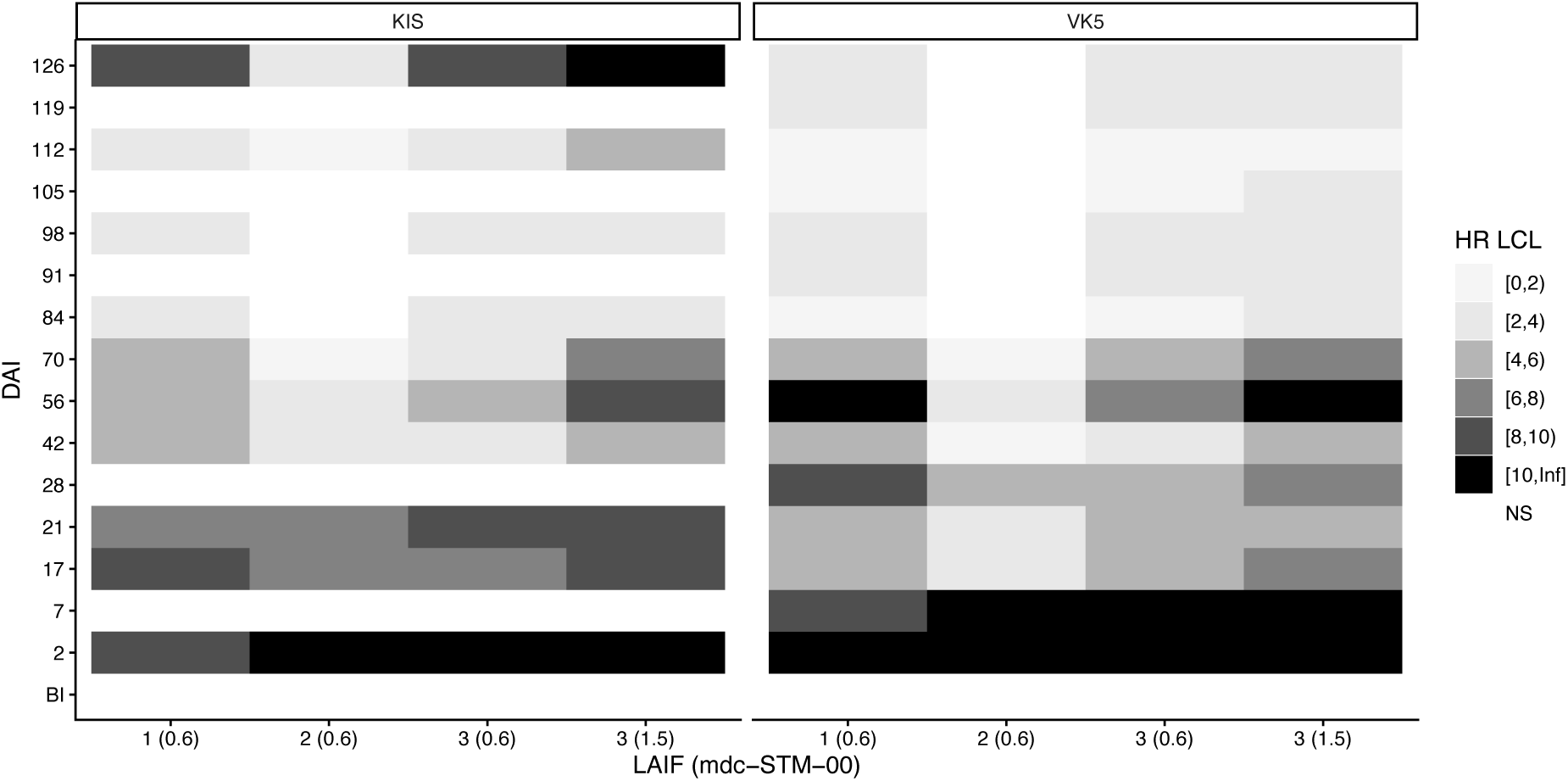
Heat maps illustrating the lower 95% confidence limit (LCL) of 10-day mortality hazard ratios (HRs) for KIS and VK5 colony mosquitoes fed at different DAIs on cattle treated with the formulations mdc-STM-001-0.6, mdc-STM-002-0.6, mdc-STM-003-0.6 or mdc-STM-003-1.5. HR and *P-values* for each DAI are available in Additional file 1: Table S7. BI is the timepoint before injection. Results for 4- and 30-day HRs are given in the Additional file 1: Fig. S4.

The mdc-STM-002-0.6 formulation was clearly the least effective, with HRs ≥ 4 for the shortest duration regardless of colony (Figure 3). It was therefore excluded from further efficacy comparisons.

The mdc-STM-001-0.6 formulation induced sustained 10-days HR ≥ 4 through 70 days post-dose for both KIS and VK5 strains. The mdc-STM-003-06 display less sustained efficacy, with HRs being superior to 4 at DAI 42 for both KIS and VK5 strains. The higher-dose of the mdc-STM-003-1.5 showed a similar pattern to mdc-STM-001-0.6, with the benefit of 10-day HRs >4 at an additional time point for KIS colony (DAI = 112).

##### Median survival times

Median survival times were consistently greater than 10 days for KIS and VK5 mosquitoes fed on control calves (Figure 4) for all the DAI tested.

**Figure 4:**
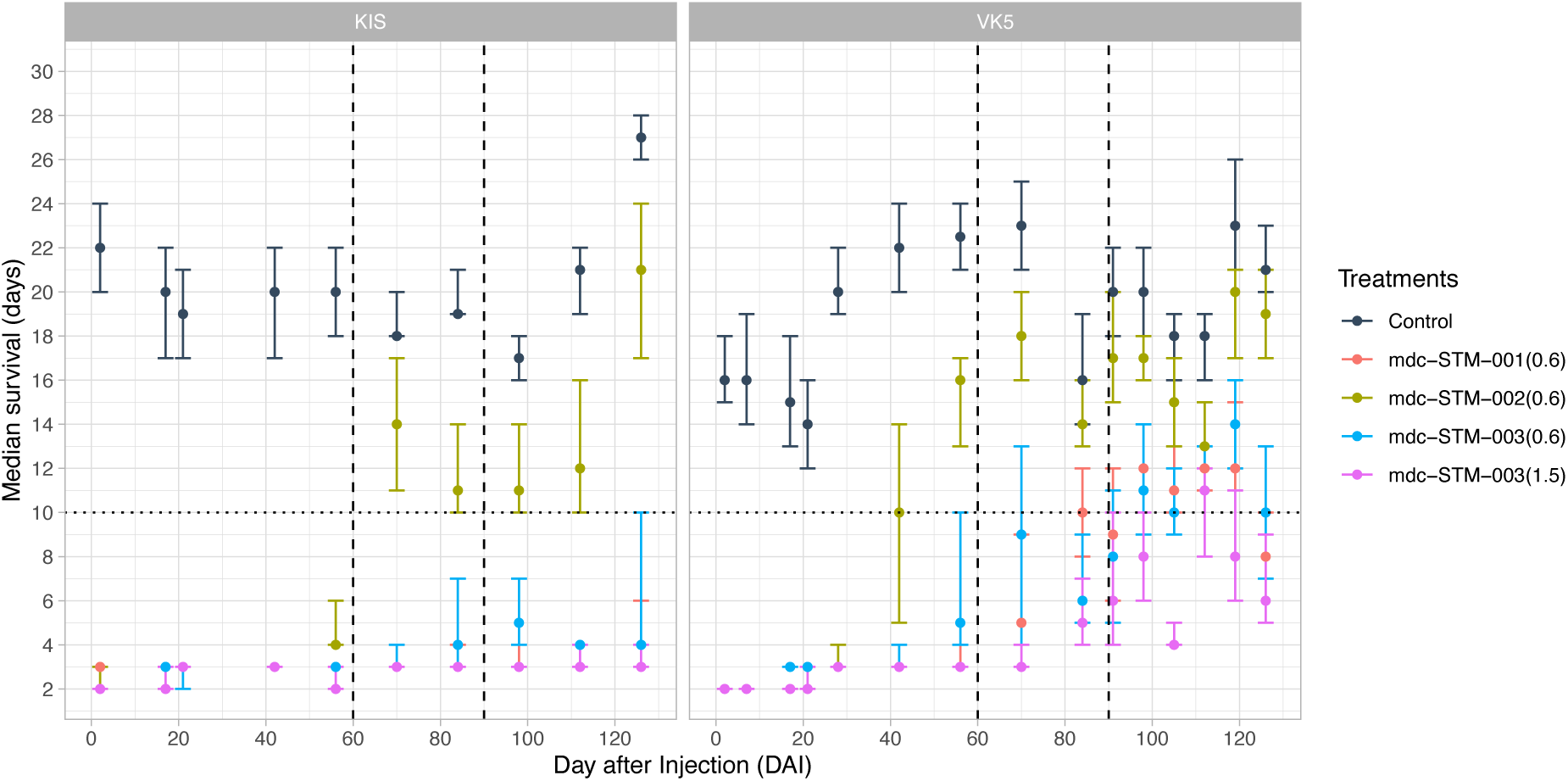
Median survival times of mosquitoes from both KIS and VK5 colonies fed at different days after injection of the LAIFs mdc-STM-001-0.6, mdc-STM-002-0.6, mdc-STM-003-06 and mdc-STM-003-1.5. The dashed horizontal line represents the average extrinsic incubation period (EIP) for *Plasmodium falciparum*. The dashed vertical lines represent 2-and 3-months post injections. Dots represent median values, +/- 95% CI are given.

For KIS mosquitoes, median survival times remained consistently below 10 days throughout the entire follow-up when fed on cattle treated with mdc-STM-001-0.6, mdc-STM-003-0.6 and mdc-STM-003-1.5. In contrast, mdc-STM-002-0.6, induced median survival times equal to or greater than the 10-day threshold from DAI 70 onward.

For VK5 mosquitoes, median survival times remained ≤ 10 days up to 91 days post-treatment, and again at DAI 126, when exposed to cattle treated with mdc-STM-001-0.6, mdc-STM-003-0.6, and mdc-STM-003-1.5 (Figure 4). The LAIF mdc-STM-002-0.6 maintained median survival times below or equal to the EIP threshold for only 40 days post-dose.

Notably, mdc-STM-001-0.6 and mdc-STM-003-1.5 induced particularly short survival durations, with median survival times <4 days for nearly 100 days in KIS mosquitoes and approximately 60 days in VK5.

##### Lethal concentrations and coverage duration

Concentration/response relationships were examined for each colony and for each LAIF. No significant differences in LC50 values were found between LAIFs, regardless of the mosquito colony (Additional file 1: Tables S9–S12; Additional file 2: Fig. S4 and S5). A unified concentration–response model, including mosquito colony as explanatory variable was thus developed using the full dataset (Figure 5).

**Figure 5.**
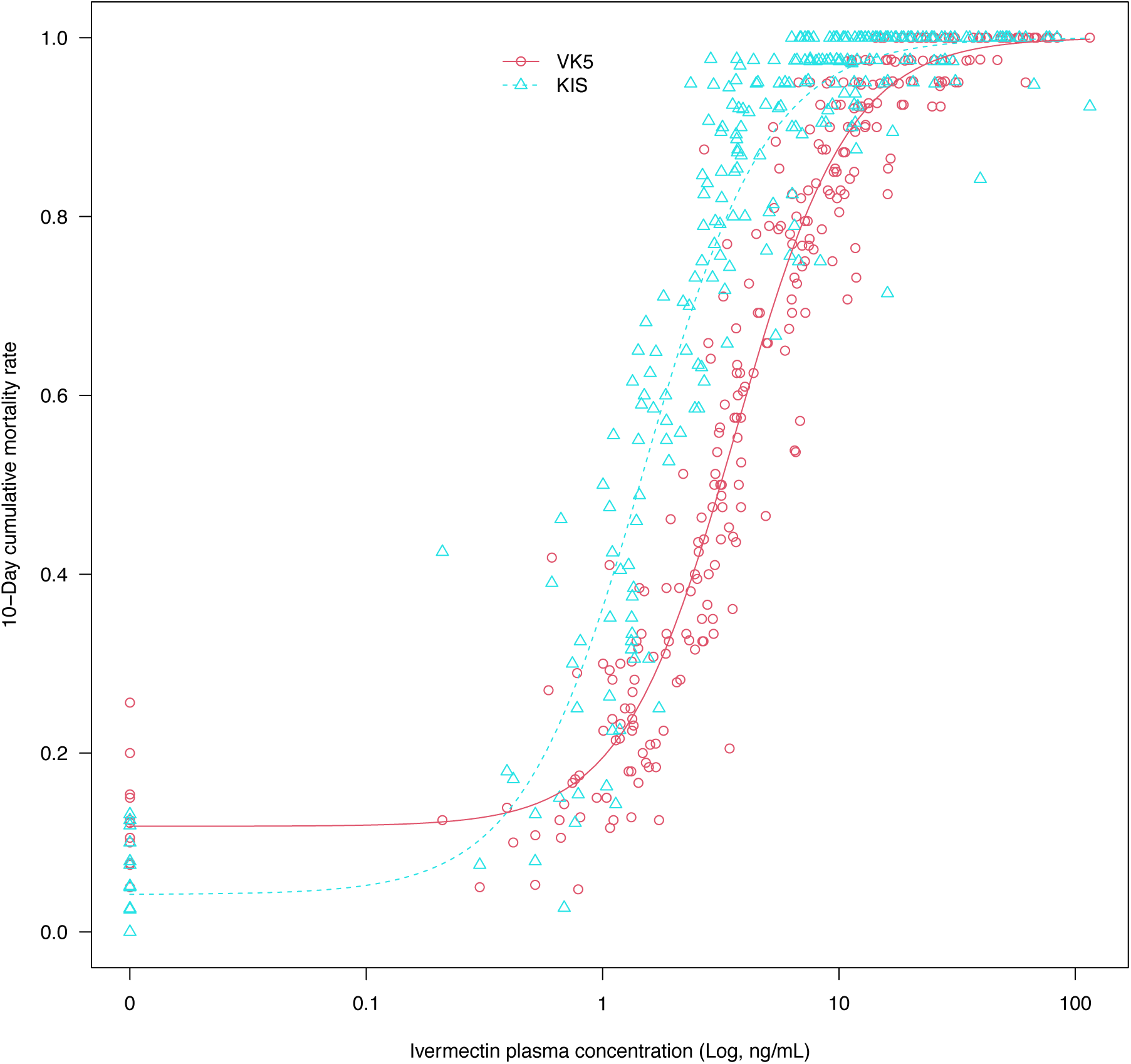
Relationship between ivermectin plasma concentrations and 10-day cumulative mosquito mortality for KIS and VK5 colonies. Data for 4-day cumulative mortality are shown in the Additional file 2: Fig. S6.

A significant difference in cumulative mortalities between KIS and VK5 was observed in relation to ivermectin concentrations, with 10-day LC50 values lower for KIS than for VK5 (mean LC50 (95% CI) for KIS = 1.51 [1.1-2.07], for VK5 = 3.66 [2.69-4.97], t-value = - 6.3187, *P*-value < 0.0001). Ten-day LC50 values sufficient to kill KIS and VK5 mosquitoes are sustained for the whole experiment duration for KIS and for more than 2 months for VK5, except for the mdc-STM-002 formulation (table 6). Values for 4-day and 10-day LC50 and LC90 are given in the Additional file1: Table S13.

**Table 6:** Time period during which the LAIF candidates reach ivermectin plasma concentration values above or equal to 10-day LC50 for KIS and VK5 colonies. Upper and lower confidence intervals (95%) are given in brackets.

| LAIF | KIS | VK5 |
| --- | --- | --- |
| mdc-STM-001-0.6 | 132 [132-132] | 132 [67-132] |
| mdc-STM-002-0.6 | 73 [65-119] | 50 [39-58] |
| mdc-STM-003-0.6 | 132 [132-132] | 128 [78-132] |
| mdc-STM-003-1.5 | 132 [132-132] | 132 [132-132] |

Inter-animal variability was also assessed, as it directly informs the coverage potential of each formulation, *i.e.* the proportion of treated animals maintaining mosquitocidal ivermectin plasma concentrations above the LC50 over time. The mdc-STM-001-0.6 achieved 100% coverage of VK5 10-day LC50 for a duration of at least 2 months post-dose (data not shown).

## Discussion

This study represents the first evaluation of candidate long-acting injectable formulations of ivermectin specifically developed for human use in malaria control. These formulations are issued from unique long lasting delivery technology and are designed to meet, with a single subcutaneous injection, the World Health Organization’s minimum target product profile for endectocides against malaria: achieving, with a single administration, at least one month of efficacy, defined by HR > 4 for *Plasmodium* vectors survival. The primary objective of this work was to identify the most promising formulation for further development, including progression to Phase 1 clinical trial. To guide formulation selection, we employed a comprehensive approach integrating multiple, complementary criteria related to mosquitocidal efficacy, pharmacokinetic characteristics, and inter-host variability, all critical for achieving effective and sustained vector control in real-world conditions and settings. The mdc-STM-001 formulation met all the predefined criteria and was selected as the optimal candidate for further evaluation in humans, based on the following key positive outcomes: i) good local and systemic tolerability during and after injection, with no abnormal variation in body weight; ii) optimal control of the initial burst, preventing systemic reactions during the early post-injection period; iii) most suitable ivermectin PK and PD profiles, achieving LC50 levels over the target duration, resulting in the highest sustained mosquito mortality across both insecticide susceptible and resistant *Anopheles* strains; iv) the lowest inter-host variability in ivermectin exposure; v) appropriate clearance beyond the predefined release duration, avoiding unnecessary systemic exposure.

In both magnitude and duration, the mosquitocidal efficacy of a single subcutaneous injection of the mdc-STM-001 formulation is compelling. It provides efficacy durations of at least 2 months, at least four times longer than those achieved with either gastric-resident or conventional oral formulations, regardless of the dosing regimen considered. A sustained reduction in mosquito survival rate lasting over two months is the factor expected to most strongly and negatively influence vectorial capacity [52]. Furthermore, by consistently shortening mosquito lifespan below the 10-day EIP, the mdc-STM-001 formulation has strong potential to dramatically reduce the proportion of infectious mosquitoes, and therefore malaria transmission, across most epidemiological contexts.

### Strengths and limitations of the experimental model for extrapolation to humans

We acknowledge the limitation of extrapolating human pharmacokinetic and pharmacodynamic parameters from data obtained following cattle treatment with the MDC-STM-001, as plasma concentration dynamics can vary substantially across species [53]. Because ivermectin is a highly lipophilic compound, body fat content, which also varies among species, represents a key determinant of its distribution within the organism [26]. Ivermectin tends to accumulate in adipose and dermal tissues surrounding capillaries [54] from which mosquitoes obtain their blood meals. This may potentially lead to higher ivermectin concentrations being ingested by mosquitoes than those measured in venous plasma and used for PK/PD analysis [55]. Indeed, a clinical study reported ivermectin concentrations to be approximately 1.33-fold higher in capillary than in venous blood [56] even though this difference did not translate into measurable differences in mosquito mortality. In addition, cattle-derived secondary metabolites may also modulate the mosquitocidal effects of long-acting formulations, as has been suggested in humans [57] with possible species-specific differences in both the nature and kinetics of these metabolites (manuscript in preparation). For all the above reasons valid whatever the route of administration of ivermectin, extrapolation of PK and PD data from cattle to humans must be approached with caution and venous plasma ivermectin concentrations should be regarded only as proxies for the actual levels of ivermectin and its metabolites ingested by mosquitoes. Nevertheless, it is noteworthy that in a phase 1 human trial using oral ivermectin, the venous plasma 10-day LC₅₀ values lethal to *An. gambiae s.s.* fell within the same range as those obtained in the present study [26,58].

Although our experimental cattle model, like any preclinical animal system, is subject to criticism regarding pharmacokinetic extrapolations to humans, it was deliberately selected to capture field relevant *Anopheles gambiae s.l.* fitness and ivermectin mosquitocidal effects since we used mosquito lots derived from a recently founded colony from Bobo Dioulasso surroundings. Indeed, throughout the Sudano-Sahelian zone, *Anopheles* mosquitoes frequently feed on domestic animals, particularly calves, which represent major blood-meal sources in areas of high bed net use [59,60]. Mosquitoes feeding on calves exhibit survival rates and median lifespans comparable to those feeding on human [61], reinforcing the biological relevance of this model. In addition, direct skin-feeding experiments on Burkinabé local calves are highly practical and result in consistently high feeding success, as these animals are natural alternative hosts for major malaria vectors. Our model therefore allows to maximize the proportion of blood-fed mosquitoes and to reflect their true post-feeding survival potential after feeding on treated or control hosts. Collectively, these parameters provide robust and field transposable meaningful entomological endpoints that are essential for predicting the performance of the long-acting ivermectin formulation in real settings.

### Mosquito resistance to ivermectin and mitigation plans

Resistance to ivermectin exists in different arthropods species [62,63]. Equally to oral formulation multiple dosing regimen, prolonged systemic exposure to ivermectin following treatment with the LAIF may exert selective pressure on resistance mechanisms, not only on its intended targets, human endoparasites, but also on *Anopheles* mosquitoes and other ectoparasites. Pyrethroids and ivermectin cross-resistance has been shown in permethrin resistant *Aedes aegypti* colonies [64]. Given the distinct molecular targets of these two compound classes, the role of metabolic resistance pathways has been hypothesized. These mechanisms often involve the overexpression of detoxification enzymes such as cytochrome P450s and xenobiotic efflux pumps [65,66]. Supporting this, dual inhibition of P450s and xenobiotic transporters was found to significantly increase ivermectin susceptibility in *An. gambiae*, highlighting detoxification pathways as a credible resistance route [67]. Ivermectin resistance in *Anopheles gambiae* could also arise due to existing alternative splicing of the gene coding for the GluCl channel, producing protein variants with differing sensitivities to ivermectin [68], which suggests that regulatory changes in splicing may contribute to reduced susceptibility.

Our findings reveal clear differences in ivermectin susceptibility between mosquito strains: the VK5 colony from *An. coluzzii* required higher plasma lethal concentrations than the fully susceptible *An. gambiae* KIS strain, suggesting the presence of tolerance mechanisms in VK5. Differences in susceptibility to ivermectin has been already shown among different *Anopheles* species [22,69,70]. In this study, we monitored the frequency of pyrethroid-resistance alleles in the VK5 mosquitoes lots, which was high throughout the experiment; however, further individual-level characterization will be needed to determine whether a direct association, if any, exists between pyrethroid-resistance genotypes and ivermectin tolerance in VK5. Beyond metabolic processes, factors such as distinct genetic backgrounds or morphological traits such as body size, may contribute to an intertwined tolerance phenotype in which multiple mechanisms interact, making it difficult to disentangle their respective roles.

The risk of selecting ivermectin resistant *Anopheles* populations during mass treatments campaigns does indeed exist, and could compromise sustained success of ivermectin-based malaria vector control strategies. In addition to physiological resistance, behavioral adaptations may further reduce the effectiveness of such interventions. Prolonged ivermectin exposure could also select for more zoophagic or opportunistic mosquito populations that preferentially feed on untreated animals or humans, thereby circumventing the drug’s effects and diminishing overall intervention impact. Nonetheless, current MDA protocols for ivermectin include exclusion criteria that prevent treatment of pregnant or lactating women and children under 15 kg. Women of child bearing potential might further be excluded in case long-acting formulation deployment. These untreated subpopulations can act as genetic refuges, reservoirs of susceptible genotypes, which help maintain genetic diversity in vector and parasite populations and reduce the overall selection pressure for resistance [40,71,72]. Such refuges could contribute to the long-term sustainability of ivermectin-based interventions.

To preserve efficacy, it is critical that implementation strategies incorporate routine surveillance of both behavioral and physiological resistance [71]. However, there is currently no established and standardized assay for detecting ivermectin resistance in mosquitoes. While molecular markers will be instrumental in tracking the emergence and spread of resistance alleles, complementary phenotypic assays are urgently needed to detect functional resistance in the field. Traditional phenotypic methods rely on mosquitoes ingesting ivermectin via membrane feeding or direct feeding on treated hosts. These approaches, however, face practical limitations, membrane feeding often results in low blood uptake by mosquitoes, and direct feeding raises ethical concerns. A promising alternative involves delivering ivermectin-laced blood using filter papers [73].

### Coverage of blood sources

The residence time of ivermectin at efficient mosquitocidal concentrations in human blood is the principal determinant of its epidemiological impact, together with the proportion of the population exposed at such concentrations [74]. This has been demonstrated using modeling studies [30,75]. The target population for standard oral ivermectin administration excludes pregnant women, nursing mothers, and children weighing under 15 kg [76,77], whereas the prolonged duration of action of mdc-STM-001 may also necessitate excluding women of childbearing potential to avoid fetal exposure. This would markedly reduce the target population size and could therefore limit overall effectiveness. However, using simulated data from theoretical formulations with constant efficacy durations of 14 to 90 days, models demonstrate that, combined with SMC, longer lasting effects could lead to significant decrease in malaria burden despite the reduced target population [78]. Further mathematical modeling, building on previous work [79], is ongoing using the present data to explore the potential of mdc-STM-001 under different implementation scenarios [80].

The specific safety concern regarding the inclusion of women of childbearing potential, primarily related to possible fetal toxicity, is debated [81], as is the inclusion of children weighing less than 15 kg [82]. Addressing these important matters will enable accurate projections for future study designs and implementation strategies.

As already mentioned above, the effectiveness of long-acting ivermectin interventions will strongly depend on achieving sufficient coverage of the blood sources available to *Anopheles* mosquitoes, both human and animal. Detailed knowledge of mosquito bionomics within potential implementation zones is therefore critical to success [36], particularly since large regions of high malaria burden correspond to the Sudano-Sahelian belt, where vectors are highly opportunistic in their feeding behaviour and often use domestic animals as alternative blood sources [59,60]. In such settings, treating livestock with endectocidal compounds has been proposed as a promising complementary strategy [83]. Commercial veterinary injectable formulations have demonstrated efficacy during up to 1-2 months against Asian vectors in buffalo and cattle, respectively [69]. A veterinary prototype long-lasting formulation from the BEPO® technology extended this efficacy over 6 months for *An. coluzzii* [40], while silicone implants achieved comparable results [84]. Beyond enhancing impact on vector populations, targeting both human and animal hosts could also contribute to mitigating the risk of ivermectin resistance if a distinct endectocide were used in animals.

### Acceptability and adherence

The deployment of a long-acting injectable ivermectin formulation for malaria vector control would represent an innovative addition to the portfolio of community-based malaria interventions. Like other public health strategies relying on high population coverage, such as vaccination programs, anti-helminthic MDAs, or malaria vector control campaigns, its success will depend on strong acceptability and adherence. Conceptually aligned with transmission-blocking vaccines (TBVs), the LAIF acts primarily through community protection. Regarding the molecule, evidence from decades of oral ivermectin MDAs for filarial control demonstrates high community acceptability, often above 90%, although adherence tends to vary across rounds [85]. The same trend is observed when ivermectin is proposed as a vector control tool against malaria [86]. Similarly, the 4 dose injectable vaccine RTS,S/AS01 have achieved strong community acceptance (80–95%) when supported by effective communication and community engagement [87,88]. Acceptability studies for TBV concepts have also reported willingness- to-receive rates above 90% in high-burden settings [89]. For the long-acting injectable ivermectin formulation, the anticipated subcutaneous injection volumes (up to a maximum of 25 µL/kg) is relatively small, suggesting minimal practical or acceptability constraints for large-scale implementation. Importantly, unlike TBVs, an LAIF ivermectin intervention could also provide direct individual health benefits by preventing or treating parasitic infections traditionally controlled by ivermectin, while simultaneously reducing malaria transmission. Additional collateral benefits from ivermectin MDAs against malaria have recently been reported, including reductions in childhood anemia [35] and decreases in jigger infestations and bedbug nuisances [90]. This dual individual and community benefit may substantially enhance perceived value, acceptability, and adherence compared to TBVs. By combining infrequent dosing, a familiar and trusted product with a well-known safety profile [91], and a clear community rationale, ivermectin MDA using LAIF could become a socially acceptable and operationally feasible complement to existing malaria control and vaccination platforms. Furthermore, such strategy could leverage the logistical infrastructure and community trust established through vaccination programs, facilitating delivery, adherence monitoring, and social acceptance.

## Conclusion

This study demonstrates that among the three long-acting injectable ivermectin formulations developed for human use, the mdc-STM-001 formulation exhibits the most favorable combination of pharmacokinetic properties, pharmacodynamic efficacy, and tolerability in a cattle-mosquito model. A single subcutaneous administration of mdc-STM-001 achieved sustained mosquitocidal concentrations over two months, inducing high mortality in both pyrethroid-susceptible and -resistant *Anopheles* strains, thereby meeting and exceeding the WHO-defined minimum criteria for endectocides targeting malaria vectors. These findings support the potential of mdc-STM-001 as a promising tool to complement existing vector control strategies, particularly in areas with behavioral or physiological resistance to conventional insecticides. While extrapolation from cattle to humans warrants caution, the observed pharmacological profile and robust entomological endpoints provide a strong rationale for advancing this formulation into Phase 1 clinical trials. Safety will remain a key consideration given the long-lasting systemic exposure; although the multiple administrations of high oral dose are safe which is supporting a favorable risk–benefit perspective for further human evaluation. The formulation will also retain the positive collateral benefits observed with oral ivermectin, such as reductions in parasitic infections, childhood anemia, and ectoparasite nuisances, underscoring the value of mdc-STM-001 as both a community- and individual-level intervention, potentially further enhancing community acceptability and adherence. Future investigations should leverage modeling approaches to optimize dosing regimens and explore synergistic integration with other vector control strategies, including seasonal malaria chemoprevention, long-lasting insecticidal nets, indoor residual spraying, or livestock-targeted endectocide interventions. Such models will also help anticipate the population coverage and logistical requirements needed to maximize epidemiological impact while ensuring operational feasibility. Given the potential for physiological or behavioral resistance to emerge, continued studies on mitigating resistance dynamics in vector populations will be essential to ensure sustainable efficacy. Overall, ivermectin MDA using the mdc-STM-001 long-acting injectable formulation based on BEPO® technology represents a potentially transformative adjunct to existing malaria control measures, with the capacity to reduce transmission by targeting both indoor- and outdoor-biting vectors over extended periods.

## Supplementary information

### Additional Tables. Additional file 1

**Table S1.** Injected volumes (mL) per animal and per formulation. **Table S2.** Median weight (Q1, Q3) in kg for each experimental cattle group at each weighing date. **Table S3**. Additional Mean (SD) pharmacokinetic parameters of ivermectin in cattle plasma. **Table S4**. Ivermectin extraction results in each depot collected at study end. **Table S5.** Copolymers extraction results in each depot collected at study end. **Table S6.** Number of mosquitoes homozygous or heterozygous for the mutated KDR-W allele. Tables S7A and S7B. Number of KIS (A) and VK5 (B) blood fed mosquitoes followed for their survival throughout the experiment. **Tables S8.** Hazard ratios (HRs), z-values, and associated p-values derived from the Cox proportional hazards models for KIS and VK5 mosquitoes. **Tables S9-S12**: between formulations comparisons of lethal concentrations. **Table S13.** Lethal concentration (LC50) for KIS and VK5 mosquitoes over 4-day follow up period (4-day LC50).

### Additional Figures. Additional file 2

**Fig. S1.** Additional mean concentration–time profiles of ivermectin in cattle plasma. **Fig. S2.** Kaplan-Meier plots. **Fig. S3.** 4-day cumulative mortality of KIS and VK5. **Fig. S4.** Heat maps illustrating the lower 95% confidence limit (LCL) of 4-day and 30-day mortality hazard ratios (HRs) for KIS and VK5 colony mosquitoes. **Fig. S5.** Relationship between ivermectin plasma concentration and 4-days (A and B) or 10-day (C and D) cumulative mortality for each candidate formulation. Fig. S6. Relationship between ivermectin plasma concentrations and 4-day cumulative mosquito mortality for KIS and VK5 colonies.

## Abbreviations

CIRDES: Centre international de recherche-développement sur l’élevage en zones subhumides;
DAI: Day after injection;
IRSS: Institut de Recherche en Sciences de la Santé;
K2-EDTA: Dipotassium Ethylenediaminetetraacetic acid;
KDR: Knock Down Resistance;
LAIF: Long acting ivermectin formulation;
LC: Lethal concentration;
LC-MS/MS: Liquid Chromatography – Tandem Mass Spectrometry;
MDA: Mass drug administration;
PD: Pharmacodynamic;
PK: Pharmacokinetic;
qPCR: Quantitative polymerase chain reaction;
RCT: Randomized controlled trial;
SMC: Seasonal malaria chemoprevention;
TBV: Transmission blocking vaccines;
WHO: World health organization

## Acknowledgements

We are particularly grateful to Boly Saïdou for his dedicated care of the animals in the stable and for his technical support during animal-related procedures. We thank Kambou Nourou Ramzi for his valuable help in mosquito husbandry and for the assistance provided during the experiments. We are also grateful to Da Firmin for his assistance with insectary husbandry, and to Da Firmin and Millogo Léopold for their technical assistance during the molecular work. We would like to thank the two anonymous reviewers for their advice and constructive comments, which greatly improved our manuscript.

## Declarations

### Availability of data and material

The datasets and code used for the analyses and figure generation reported in this study are publicly available at https://doi.org/10.23708/PSJMLM

### Authors’ contribution

LZ: Investigation, preliminary statistical analysis, writing original draft. SHP, KM, NM, CP, KRD, SLD: Methodology and protocol design. SHP: Supervision and coordination of experimental work at CIRDES, conceptualisation. COW: Investigation, editing. AFS: Supervision of COW, editing. AS: Molecular biology coordination and writing, editing. AP: Formal analysis, data curation, and validation. SB: Visualization. SLD: Pharmacokinetic coordination, writing, and review. KM: Supervision, conceptualization, funding acquisition, administrative management, and writing, review and editing. CR: Conceptualization and funding acquisition. AMGB: Supervision and mentoring of LZ (thesis director). G-KD: Resources and administrative management. All authors read and approved the final manuscript.

### Ethics approval

This study is part of IMPACT project that got approval from the CIRDES ethics committee under letter no. 004-10/2020 CEB-CIRDES. The study involving animals has been conducted ethically following the governance of animal care and use for scientific purposes in Africa and the Middle East [92].

### Consent for publication

Not applicable

## Supplementary tables

**Supplementary Table S1.** Injected volumes (mL) per animal and per formulation, given the body weight (BW) measured the day of the treatment (kg)

| Animal ID | BW at arrival (kg) | BW at injection (kg) | Formulation ID | Injected volume (mL) |
| --- | --- | --- | --- | --- |
| B122 | 141.6 | 153.4 | mdc-STM-001 | 1.18 |
| B126 | 116.8 | 140.2 | mdc-STM-001 | 1.08 |
| B128 | 109.4 | 119.8 | mdc-STM-001 | 0.92 |
| B332 | 119.0 | 126.6 | mdc-STM-001 | 0.98 |
| B335 | 99.2 | 115.2 | mdc-STM-001 | 0.89 |
| B121 | 107.2 | 120.8 | mdc-STM-002 | 0.94 |
| B130 | 128.8 | 137.6 | mdc-STM-002 | 1.07 |
| B324 | 97.4 | 115.2 | mdc-STM-002 | 0.90 |
| B330 | 138.0 | 144.6 | mdc-STM-002 | 1.13 |
| B334 | 142.0 | 151.4 | mdc-STM-002 | 1.18 |
| B123 | 101.2 | 122.3 | mdc-STM-003 | 0.72 |
| B124 | 124.4 | 130.6 | mdc-STM-003 | 0.76 |
| B125 | 124.6 | 142.2 | mdc-STM-003 | 0.83 |
| B326 | 140.8 | 160.8 | mdc-STM-003 | 0.94 |
| B333 | 94.2 | 106.6 | mdc-STM-003 | 0.62 |
| B322 | 127.4 | 146.8 | mdc-STM-003 | 2.14 |
| B323 | 108.6 | 120.0 | mdc-STM-003 | 1.75 |
| B327 | 129.4 | 144.6 | mdc-STM-003 | 2.11 |
| B329 | 99.8 | 115.8 | mdc-STM-003 | 1.69 |
| B331 | 106.4 | 124.8 | mdc-STM-003 | 1.82 |

**Supplementary Table S2.**
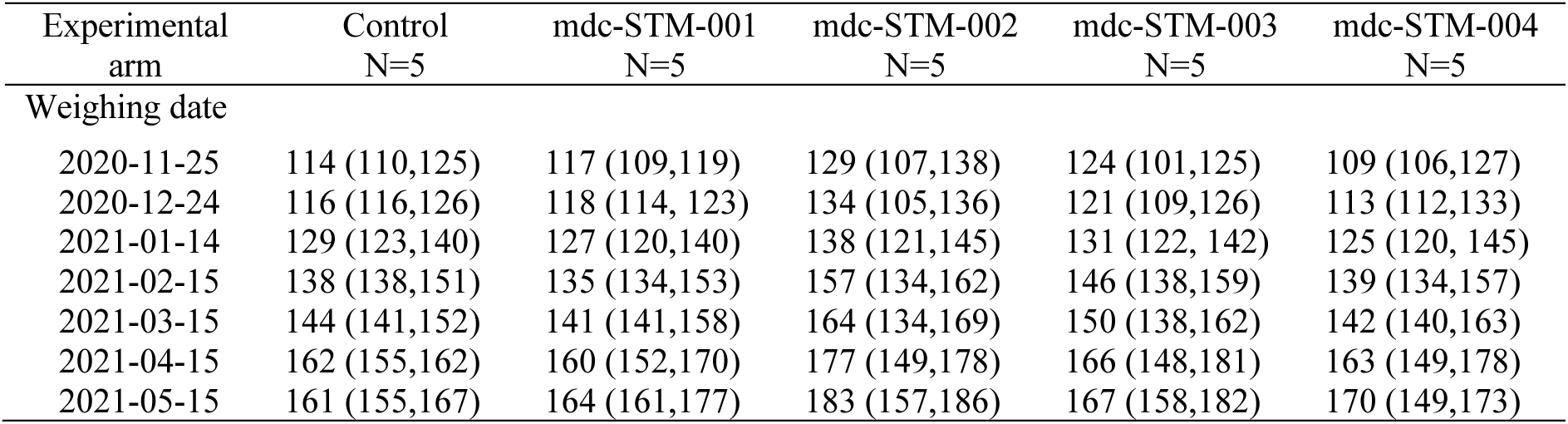
Median weight (Q1, Q3) in kg for each experimental cattle group at each weighing date.

| Experimental<br>arm | Control<br>N=5 | mdc-STM-001<br>N=5 | mdc-STM-002<br>N=5 | mdc-STM-003<br>N=5 | mdc-STM-004<br>N=5 |
| --- | --- | --- | --- | --- | --- |
| Weighing date |  |  |  |  |  |
| 2020-11-25 | 114 (110,125) | 117 (109,119) | 129 (107,138) | 124 (101,125) | 109 (106,127) |
| 2020-12-24 | 116 (116,126) | 118 (114, 123) | 134 (105,136) | 121 (109,126) | 113 (112,133) |
| 2021-01-14 | 129 (123,140) | 127 (120,140) | 138 (121,145) | 131 (122, 142) | 125 (120, 145) |
| 2021-02-15 | 138 (138,151) | 135 (134,153) | 157 (134,162) | 146 (138,159) | 139 (134,157) |
| 2021-03-15 | 144 (141,152) | 141 (141,158) | 164 (134,169) | 150 (138,162) | 142 (140,163) |
| 2021-04-15 | 162 (155,162) | 160 (152,170) | 177 (149,178) | 166 (148,181) | 163 (149,178) |
| 2021-05-15 | 161 (155,167) | 164 (161,177) | 183 (157,186) | 167 (158,182) | 170 (149,173) |

**Supplementary Table S3.** Mean (SD) pharmacokinetic parameters of ivermectin in cattle plasma after single subcutaneous administration of LAIFs at 0.6 or 1.5 mg/kg.

| Parameters | mdc-STM-001-0.6 | mdc-STM-002-0.6 | mdc-STM-003-0.6 | mdc-STM-003-1.5 |
| --- | --- | --- | --- | --- |
| Tmax<sup>a</sup> (h) | 24 (24; 48) | 48 (24; 168) | 24 (24; 48) | 24 (24; 168) |
| Cmax (ng/mL) | 35.4 (12.7) | 50.3 (13.7) | 58.1 (25.2) | 81.2 (32.2) |
| Cmax/Dose (ng/mL/mg/kg) | 58.3 (22.0) | 84.3 (22.6) | 99.4 (42.2) | 56.1 (22.4) |
| Clast (ng/mL) | 3.93 (2.75) | 1.03 (0.70) | 3.43 (2.74) | 6.30 (4.35) |
| AUC(0-28d) (h.ng/mL) | 14227 (3853) | 16253 (5465) | 18085 (9789) | 26248 (5534) |
| AUC(0-28d)/Dose (h.ng/mL/mg/kg) | 23378 (6862) | 27298 (9171) | 30956 (16689) | 18116 (3822) |
| AUC(0-91d) (h.ng/mL) | 25055 (2936) | 21121 (8866) | 28890 (18668) | 41820 (13286) |
| AUC(0-91d)/Dose (h.ng/mL/mg/kg) | 40990 (5664) | 35479 (14839) | 49483 (31994) | 28903 (9300) |
| AUClast (h.ng/mL) | 28534 (2316) | 22322 (9059) | 32851 (21691) | 47192 (15724) |
| AUClast/Dose (h.ng/mL/mg/kg) | 46589 (3960) | 37497 (15165) | 56269 (37147) | 32630 (11042) |
| T1/2 (h) | 784 (244) (n=4) | 731 (579) (n=4) | 728 (399) (n=3) | 489 (n=1) |
| C28d (ng/mL) | 13.7 (1.6) | 8.90 (4.85) | 11.1 (8.3) | 15.5 (5.1) |
| C56d (ng/mL) | 6.70 (1.67) | 2.92 (2.63) | 7.34 (7.14) | 10.3 (6.9) |
| C91d (ng/mL) | 3.48 (1.85) | 1.11 (0.84) | 4.33 (3.99) | 5.55 (3.72) |

SD: standard deviation.
<sup>a</sup> Median (min; max).
Tmax: time to reach Cmax. Cmax: maximal plasma concentration. Clast: last quantifiable plasma concentration.
AUC(0-28d): area under curve from time 0 to 28 days. AUC(0-28d)/Dose: dose-normalized AUC(0-28d).
AUC(0-91d): area under curve from time 0 to 91 days. AUC(0-91d)/Dose: dose-normalized AUC(0-91d).
AUClast: area under curve from time 0 to last quantifiable plasma level. AUClast/Dose: dose-normalized
AUClast. T1/2: apparent terminal phase half-life. C28d: plasma concentration 28 days post injection. C56d:
plasma concentration 56 days post injection. C91d: plasma concentration 91 days post injection.

**Supplementary Table S4.** Ivermectin extraction results in each depot collected at study end.

| Treatment<br>Dose<br>(mg/kg) | Animal ID | IVM injected<br>dose (mg) | IVM recovered dose (mg) |  | IVM recovered dose (%) |  |
| --- | --- | --- | --- | --- | --- | --- |
|  |  |  | Indiv. | Mean (SD) | Indiv. | Mean (SD) |
| mdc-STM-001-0.6 | B122 | 100.4 | 38.6 |  | 38.5 |  |
|  | B126 | 85.0 | 18.3 |  | 21.6 |  |
|  | B128 | 73.3 | 2.5 | 19.4 (12.9) | 3.3 | 23.0 (12.7) |
|  | B332 | 76.3 | 20.6 |  | 27.0 |  |
|  | B335 | 67.9 | 16.8 |  | 24.8 |  |
| mdc-STM-002-0.6 | B121 | 76.1 | 39.0 |  | 51.3 |  |
|  | B130 | 84.7 | 6.1 |  | 7.2 |  |
|  | B324 | 69.0 | 23.4 | 32.3 (17.7) | 33.9 | 39.1 (19.6) |
|  | B330 | 87.7 | 41.7 |  | 47.6 |  |
|  | B334 | 92.0 | 51.1 |  | 55.5 |  |
| mdc-STM-003-0.6 | B123 | 70.6 | 5.4 |  | 7.7 |  |
|  | B124 | 76.4 | 14.3 |  | 18.7 |  |
|  | B125 | 81.1 | 42.6 | 17.8 (14.4) | 52.6 | 23.1 (17.6) |
|  | B326 | 96.4 | 12.0 |  | 12.5 |  |
|  | B333 | 61.6 | 14.8 |  | 24.0 |  |
| mdc-STM-003-1.5 | 127.4 | 211.6 | 69.8 |  | 33.0 |  |
|  | 108.6 | 173.8 | 55.5 |  | 32.0 |  |
|  | 129.4 | 212.3 | 0.5 | 59.5 (35.5) | 0.2 | 32.7 (20.0) |
|  | 99.8 | 164.8 | 79.7 |  | 48.4 |  |
|  | 106.4 | 182.9 | 91.8 |  | 50.2 |  |

IVM: ivermectin. Indiv.: individual. SD: standard deviation.

**Supplementary Table S5.** Copolymers extraction results in each depot collected at study end.

| Treatment<br>Dose (mg/kg) | Animal ID | Copolymer<br>injected (mg) | Copolymer<br>recovered (mg) | Copolymer recovered<br>(%) |  |
| --- | --- | --- | --- | --- | --- |
|  |  |  |  | Indiv. | Mean (SD) |
| mdc-STM-001-0.6 | B122 | 442.2 | 120.1 | 27.2 | 18.8 (8.5) |
|  | B126 | 374.4 | 90.0 | 24.0 |  |
|  | B128 | 323.1 | 17.3 | 5.4 |  |
|  | B332 | 336.0 | 55.5 | 16.5 |  |
|  | B335 | 299.1 | 63.3 | 21.1 |  |
| mdc-STM-002-0.6 | B121 | 217.6 | 105.8 | 48.6 | 34.1 (15.9) |
|  | B130 | 242.2 | 19.5 | 8.0 |  |
|  | B324 | 197.4 | 77.3 | 39.2 |  |
|  | B330 | 250.8 | 108.8 | 43.4 |  |
|  | B334 | 263.2 | 81.6 | 31.0 |  |
| mdc-STM-003-0.6 | B123 | 155.6 | 25.0 | 16.1 | 25.9 (25.0) |
|  | B124 | 168.4 | 11.4 | 6.8 |  |
|  | B125 | 178.6 | 39.5 | 22.1 |  |
|  | B326 | 212.4 | 32.0 | 15.1 |  |
|  | B333 | 135.6 | 94.2 | 69.5 |  |
| mdc-STM-003-1.5 | 127.4 | 466.2 | 196.4 | 42.1 | 34.1 (18.8) |
|  | 108.6 | 382.8 | 164.3 | 42.9 |  |
|  | 129.4 | 467.6 | 3.5 | 0.8 |  |
|  | 99.8 | 363.0 | 167.4 | 46.1 |  |
|  | 106.4 | 403.0 | 155.6 | 38.6 |  |

Indiv.: individual. SD: standard deviation.

**Supplementary Table S6.** Number of mosquitoes homozygous or heterozygous for the mutated KDR-W allele (resistant allele, R) of the voltage-gated sodium channel gene, which confers resistance to pyrethroids. In contrast, the wild-type allele is designated S (susceptible allele). RR and RS indicate homozygous and heterozygous genotypes for the KDR-W mutation, respectively. SS corresponds to homozygosity for the wild-type allele. Resistant phenotypes (RR and RS genotypes) among the tested mosquitoes (n) are given as percentages. Numbers are reported per generation (F2 to F9) and per day after injection (DAI) at which mosquitoes were exposed to cattle.

| DAI | Generation (F) | RR (n) | RS (n) | SS (n) | Resistant phenotypes (%) | Moquitoes tested (n) |
| --- | --- | --- | --- | --- | --- | --- |
| pre-dose | F2 | 10 | 15 | 8 | 73,52941176 | 33 |
| 2, 7, 17, 21, 28 | F3 | 2 | 6 | 2 | 80 | 10 |
| 42 | F5 | 12 | 20 | 9 | 74,41860465 | 41 |
| 56 | F6 |  |  |  | na | na |
| 70, 84, 91, 98 | F7 | 6 | 19 | 21 | 54,34782609 | 47 |
| 105,112, 119 | F8 |  |  |  | na | na |
| 126 | F9 | 5 | 14 | 17 | 52,77777778 | 36 |

**Supplementary Table S7A.** Number of KIS blood fed mosquitoes followed for their survival before injection (BI) and at the different days after injection (DAI) for each cattle of the different treatment groups.

| Mosquito Colony | Treatment | Cattle ID | BI | Day after injection (DAI) |  |  |  |  |  |  |  |  |  |  |  |  |  |  |
| --- | --- | --- | --- | --- | --- | --- | --- | --- | --- | --- | --- | --- | --- | --- | --- | --- | --- | --- |
|  |  |  |  | 2 | 7 | 17 | 21 | 28 | 42 | 56 | 70 | 84 | 91 | 98 | 105 | 112 | 119 | 126 |
| kis | Control | B127 | 40 | 31 | 0 | 37 | 39 | 0 | 19 | 29 | 41 | 40 | 0 | 40 | 37 | 38 | 39 | 39 |
|  |  | B129 | 40 | 34 | 0 | 31 | 41 | 0 | 38 | 31 | 41 | 41 | 0 | 36 | 32 | 38 | 40 | 40 |
|  |  | B321 | 40 | 34 | 0 | 38 | 34 | 0 | 39 | 38 | 41 | 40 | 0 | 0 | 0 | 39 | 43 | 41 |
|  |  | B328 | 41 | 20 | 0 | 35 | 37 | 0 | 32 | 38 | 42 | 40 | 0 | 42 | 42 | 41 | 41 | 38 |
|  |  | B336 | 38 | 34 | 0 | 37 | 42 | 0 | 19 | 32 | 41 | 40 | 0 | 39 | 39 | 41 | 39 | 40 |
|  | mdc-STM-001-0.6 | B122 | 40 | 38 | 0 | 39 | 40 | 20 | 40 | 37 | 40 | 39 | 0 | 40 | 37 | 40 | 41 | 42 |
|  |  | B126 | 39 | 40 | 0 | 40 | 40 | 21 | 40 | 38 | 38 | 36 | 0 | 40 | 40 | 42 | 38 | 36 |
|  |  | B128 | 38 | 28 | 0 | 39 | 41 | 9 | 40 | 41 | 41 | 40 | 0 | 0 | 0 | 40 | 37 | 41 |
|  |  | B332 | 40 | 41 | 0 | 34 | 39 | 21 | 38 | 42 | 40 | 44 | 0 | 38 | 40 | 43 | 39 | 38 |
|  |  | B335 | 40 | 38 | 0 | 40 | 40 | 24 | 32 | 39 | 39 | 43 | 0 | 39 | 41 | 41 | 40 | 41 |
|  | mdc-STM-002-0.6 | B121 | 38 | 38 | 0 | 40 | 43 | 10 | 37 | 44 | 38 | 40 | 0 | 39 | 36 | 41 | 40 | 40 |
|  |  | B130 | 40 | 39 | 0 | 40 | 40 | 21 | 40 | 42 | 32 | 40 | 0 | 38 | 15 | 39 | 38 | 39 |
|  |  | B324 | 40 | 39 | 0 | 40 | 40 | 21 | 40 | 39 | 40 | 40 | 0 | 37 | 40 | 36 | 39 | 37 |
|  |  | B330 | 40 | 41 | 0 | 41 | 40 | 24 | 31 | 40 | 39 | 40 | 0 | 40 | 38 | 39 | 39 | 39 |
|  |  | B334 | 40 | 38 | 0 | 40 | 40 | 18 | 40 | 38 | 39 | 43 | 0 | 0 | 0 | 42 | 40 | 38 |
|  | mdc-STM-003-0.6 | B123 | 40 | 40 | 0 | 41 | 38 | 10 | 40 | 40 | 40 | 39 | 0 | 38 | 37 | 40 | 39 | 38 |
|  |  | B124 | 38 | 39 | 0 | 37 | 39 | 6 | 40 | 40 | 40 | 42 | 0 | 39 | 42 | 38 | 37 | 40 |
|  |  | B125 | 40 | 40 | 0 | 38 | 40 | 25 | 39 | 38 | 40 | 41 | 0 | 0 | 0 | 38 | 39 | 41 |
|  |  | B326 | 40 | 37 | 0 | 39 | 40 | 16 | 40 | 38 | 40 | 42 | 0 | 39 | 37 | 39 | 40 | 40 |
|  |  | B333 | 40 | 42 | 0 | 39 | 39 | 4 | 41 | 36 | 43 | 38 | 0 | 40 | 33 | 40 | 40 | 42 |
|  | mdc-STM-003-1.5 | B322 | 40 | 39 | 0 | 39 | 40 | 20 | 40 | 39 | 40 | 42 | 0 | 41 | 37 | 38 | 42 | 40 |
|  |  | B323 | 40 | 40 | 0 | 39 | 40 | 18 | 39 | 32 | 40 | 40 | 0 | 25 | 37 | 40 | 42 | 44 |
|  |  | B327 | 40 | 35 | 0 | 38 | 40 | 17 | 41 | 24 | 38 | 41 | 0 | 43 | 26 | 36 | 39 | 41 |
|  |  | B329 | 38 | 38 | 0 | 40 | 40 | 0 | 35 | 40 | 41 | 42 | 0 | 41 | 36 | 40 | 41 | 39 |
|  |  | B331 | 40 | 38 | 0 | 38 | 40 | 6 | 30 | 39 | 39 | 40 | 0 | 0 | 0 | 42 | 38 | 39 |
| Total |  |  | 990 | 921 | 0 | 959 | 992 | 311 | 910 | 934 | 993 | 1013 | 0 | 774 | 722 | 991 | 990 | 993 |

**Supplementary Table S7B.**
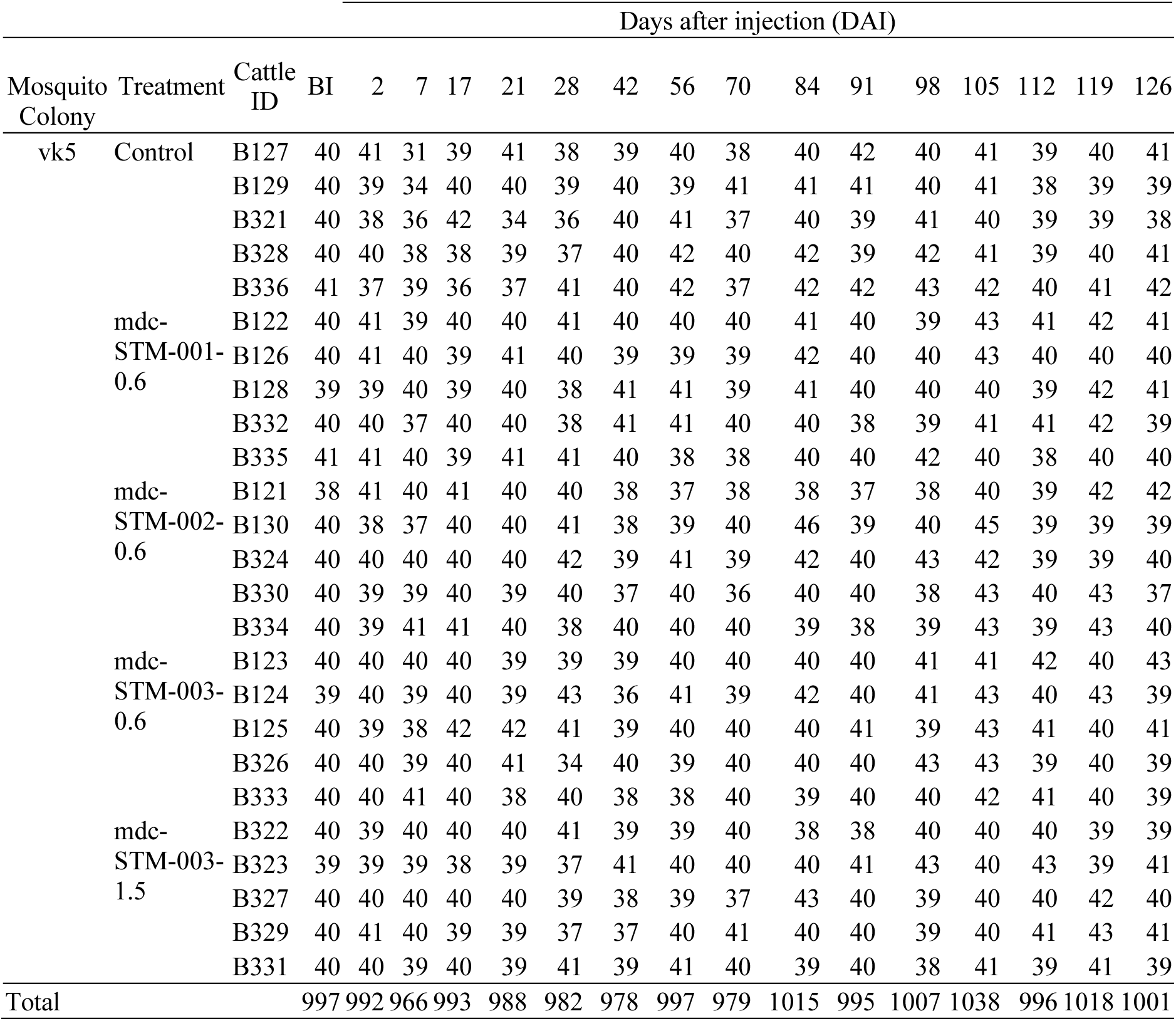
Number of VK5 blood fed mosquitoes followed for their survival before injection (BI) and at the different days after injection (DAI) for each cattle of the different treatment groups.

|  |  |  | Days after injection (DAI) |  |  |  |  |  |  |  |  |  |  |  |  |  |  |  |
| --- | --- | --- | --- | --- | --- | --- | --- | --- | --- | --- | --- | --- | --- | --- | --- | --- | --- | --- |
| Mosquito Colony | Treatment | Cattle ID | BI | 2 | 7 | 17 | 21 | 28 | 42 | 56 | 70 | 84 | 91 | 98 | 105 | 112 | 119 | 126 |
| vk5 | Control | B127 | 40 | 41 | 31 | 39 | 41 | 38 | 39 | 40 | 38 | 40 | 42 | 40 | 41 | 39 | 40 | 41 |
|  |  | B129 | 40 | 39 | 34 | 40 | 40 | 39 | 40 | 39 | 41 | 41 | 41 | 40 | 41 | 38 | 39 | 39 |
|  |  | B321 | 40 | 38 | 36 | 42 | 34 | 36 | 40 | 41 | 37 | 40 | 39 | 41 | 40 | 39 | 39 | 38 |
|  |  | B328 | 40 | 40 | 38 | 38 | 39 | 37 | 40 | 42 | 40 | 42 | 39 | 42 | 41 | 39 | 40 | 41 |
|  |  | B336 | 41 | 37 | 39 | 36 | 37 | 41 | 40 | 42 | 37 | 42 | 42 | 43 | 42 | 40 | 41 | 42 |
|  | mdc-STM-001-0.6 | B122 | 40 | 41 | 39 | 40 | 40 | 41 | 40 | 40 | 40 | 41 | 40 | 39 | 43 | 41 | 42 | 41 |
|  |  | B126 | 40 | 41 | 40 | 39 | 41 | 40 | 39 | 39 | 39 | 42 | 40 | 40 | 43 | 40 | 40 | 40 |
|  |  | B128 | 39 | 39 | 40 | 39 | 40 | 38 | 41 | 41 | 39 | 41 | 40 | 40 | 40 | 39 | 42 | 41 |
|  |  | B332 | 40 | 40 | 37 | 40 | 40 | 38 | 41 | 41 | 40 | 40 | 38 | 39 | 41 | 41 | 42 | 39 |
|  |  | B335 | 41 | 41 | 40 | 39 | 41 | 41 | 40 | 38 | 38 | 40 | 40 | 42 | 40 | 38 | 40 | 40 |
|  | mdc-STM-002-0.6 | B121 | 38 | 41 | 40 | 41 | 40 | 40 | 38 | 37 | 38 | 38 | 37 | 38 | 40 | 39 | 42 | 42 |
|  |  | B130 | 40 | 38 | 37 | 40 | 40 | 41 | 38 | 39 | 40 | 46 | 39 | 40 | 45 | 39 | 39 | 39 |
|  |  | B324 | 40 | 40 | 40 | 40 | 40 | 42 | 39 | 41 | 39 | 42 | 40 | 43 | 42 | 39 | 39 | 40 |
|  |  | B330 | 40 | 39 | 39 | 40 | 39 | 40 | 37 | 40 | 36 | 40 | 40 | 38 | 43 | 40 | 43 | 37 |
|  |  | B334 | 40 | 39 | 41 | 41 | 40 | 38 | 40 | 40 | 40 | 39 | 38 | 39 | 43 | 39 | 43 | 40 |
|  | mdc-STM-003-0.6 | B123 | 40 | 40 | 40 | 40 | 39 | 39 | 39 | 40 | 40 | 40 | 40 | 41 | 41 | 42 | 40 | 43 |
|  |  | B124 | 39 | 40 | 39 | 40 | 39 | 43 | 36 | 41 | 39 | 42 | 40 | 41 | 43 | 40 | 43 | 39 |
|  |  | B125 | 40 | 39 | 38 | 42 | 42 | 41 | 39 | 40 | 40 | 40 | 41 | 39 | 43 | 41 | 40 | 41 |
|  |  | B326 | 40 | 40 | 39 | 40 | 41 | 34 | 40 | 39 | 40 | 40 | 40 | 43 | 43 | 39 | 40 | 39 |
|  |  | B333 | 40 | 40 | 41 | 40 | 38 | 40 | 38 | 38 | 40 | 39 | 40 | 40 | 42 | 41 | 40 | 39 |
|  | mdc-STM-003-1.5 | B322 | 40 | 39 | 40 | 40 | 40 | 41 | 39 | 39 | 40 | 38 | 38 | 40 | 40 | 40 | 39 | 39 |
|  |  | B323 | 39 | 39 | 39 | 38 | 39 | 37 | 41 | 40 | 40 | 40 | 41 | 43 | 40 | 43 | 39 | 41 |
|  |  | B327 | 40 | 40 | 40 | 40 | 40 | 39 | 38 | 39 | 37 | 43 | 40 | 39 | 40 | 40 | 42 | 40 |
|  |  | B329 | 40 | 41 | 40 | 39 | 39 | 37 | 37 | 40 | 41 | 40 | 40 | 39 | 40 | 41 | 43 | 41 |
|  |  | B331 | 40 | 40 | 39 | 40 | 39 | 41 | 39 | 41 | 40 | 39 | 40 | 38 | 41 | 39 | 41 | 39 |
| Total |  |  | 997 | 992 | 966 | 993 | 988 | 982 | 978 | 997 | 979 | 1015 | 995 | 1007 | 1038 | 996 | 1018 | 1001 |

**Supplementary table S8.** Hazard ratios (HRs), z-values, and associated p-values derived from the Cox proportional hazards models. Values are given for 4, 10 and 30-days of survival follow-up after mosquitoes blood feeding, before injection (BI) and for each day after injection (DAI). A. KIS mosquitoes. B. VK5 mosquitoes.

| Formulation | Treatment arm | DAI | Follow-up<br>(days) | HR (CI 95%) | z.ratio | p.value |
| --- | --- | --- | --- | --- | --- | --- |
| (MDC-STM-001-0.6) | Control | BI | 4 | 1 (0.1-17.2) | 0.0018088 | 1.0000000 |
| (MDC-STM-002-0.6) | Control | BI | 4 | 1.9 (0.2-22.7) | 0.6900508 | 0.9587058 |
| (MDC-STM-003-0.6) | Control | BI | 4 | 0.8 (0-14.3) | -0.1754490 | 0.9997884 |
| (MDC-STM-003-1.5) | Control | BI | 4 | 1.9 (0.2-22.4) | 0.6751535 | 0.9618095 |
| (MDC-STM-001-0.6) | Control | 2 | 4 | 32.2 (9.8-106.3) | 7.9378105 | 0.0000000 |
| (MDC-STM-002-0.6) | Control | 2 | 4 | 59.3 (18-195.1) | 9.3486912 | 0.0000000 |
| (MDC-STM-003-0.6) | Control | 2 | 4 | 58.3 (17.7-191.8) | 9.3098183 | 0.0000000 |
| (MDC-STM-003-1.5) | Control | 2 | 4 | 57.3 (17.4-188.9) | 9.2598711 | 0.0000000 |
| (MDC-STM-001-0.6) | Control | 17 | 4 | 56 (17-184.3) | 9.2185509 | 0.0000000 |
| (MDC-STM-002-0.6) | Control | 17 | 4 | 53.4 (16.2-175.7) | 9.1132259 | 0.0000000 |
| (MDC-STM-003-0.6) | Control | 17 | 4 | 50.4 (15.3-165.8) | 8.9748191 | 0.0000000 |
| (MDC-STM-003-1.5) | Control | 17 | 4 | 60.6 (18.4-199.5) | 9.3992086 | 0.0000000 |
| (MDC-STM-001-0.6) | Control | 21 | 4 | 25.5 (9.3-70.3) | 8.7248194 | 0.0000000 |
| (MDC-STM-002-0.6) | Control | 21 | 4 | 23.4 (8.5-64.4) | 8.4758112 | 0.0000000 |
| (MDC-STM-003-0.6) | Control | 21 | 4 | 27.7 (10.1-76.4) | 8.9365409 | 0.0000000 |
| (MDC-STM-003-1.5) | Control | 21 | 4 | 27.7 (10-76.2) | 8.9388090 | 0.0000000 |
| (MDC-STM-001-0.6) | Control | 42 | 4 | 41.1 (10.2-165.9) | 7.2625835 | 0.0000000 |
| (MDC-STM-002-0.6) | Control | 42 | 4 | 24 (5.9-97.5) | 6.1919836 | 0.0000000 |
| (MDC-STM-003-0.6) | Control | 42 | 4 | 28.2 (7-114.1) | 6.5248654 | 0.0000000 |
| (MDC-STM-003-1.5) | Control | 42 | 4 | 40.3 (10-162.9) | 7.2196770 | 0.0000000 |
| (MDC-STM-001-0.6) | Control | 56 | 4 | 55.9 (14-223.3) | 7.9184506 | 0.0000000 |
| (MDC-STM-002-0.6) | Control | 56 | 4 | 23.6 (5.8-95) | 6.1826656 | 0.0000000 |
| (MDC-STM-003-0.6) | Control | 56 | 4 | 52.2 (13-209.7) | 7.7629011 | 0.0000000 |
| (MDC-STM-003-1.5) | Control | 56 | 4 | 81.4 (20.3-326.2) | 8.6502114 | 0.0000000 |
| (MDC-STM-001-0.6) | Control | 70 | 4 | 26.4 (8.2-84.9) | 7.6620884 | 0.0000000 |
| (MDC-STM-002-0.6) | Control | 70 | 4 | 6.2 (1.9-20.7) | 4.1257303 | 0.0003562 |
| (MDC-STM-003-0.6) | Control | 70 | 4 | 19.8 (6.1-64) | 6.9522992 | 0.0000000 |
| (MDC-STM-003-1.5) | Control | 70 | 4 | 34.4 (10.7-110.5) | 8.2674788 | 0.0000000 |
| (MDC-STM-001-0.6) | Control | 84 | 4 | 7.9 (2.9-21.6) | 5.5987636 | 0.0000002 |
| (MDC-STM-002-0.6) | Control | 84 | 4 | 2.2 (0.8-6.2) | 1.9916567 | 0.2699699 |
| (MDC-STM-003-0.6) | Control | 84 | 4 | 6.8 (2.5-18.8) | 5.1726904 | 0.0000023 |
| (MDC-STM-003-1.5) | Control | 84 | 4 | 12.8 (4.7-34.9) | 6.9245385 | 0.0000000 |
| (MDC-STM-001-0.6) | Control | 98 | 4 | 38.4 (8.7-169.4) | 6.7085831 | 0.0000000 |
| (MDC-STM-002-0.6) | Control | 98 | 4 | 2.7 (0.5-13.6) | 1.6602615 | 0.4587964 |
| (MDC-STM-003-0.6) | Control | 98 | 4 | 21.2 (4.7-94.6) | 5.5628717 | 0.0000003 |
| (MDC-STM-003-1.5) | Control | 98 | 4 | 36.7 (8.3-162.4) | 6.6104282 | 0.0000000 |
| (MDC-STM-001-0.6) | Control | 112 | 4 | 25.7 (6.4-103.6) | 6.3494193 | 0.0000000 |
| (MDC-STM-002-0.6) | Control | 112 | 4 | 5.6 (1.3-23.6) | 3.2765643 | 0.0092896 |
| (MDC-STM-003-0.6) | Control | 112 | 4 | 23.5 (5.8-95.2) | 6.1673392 | 0.0000000 |
| (MDC-STM-003-1.5) | Control | 112 | 4 | 26.8 (6.6-108) | 6.4270674 | 0.0000000 |
| (MDC-STM-001-0.6) | Control | 126 | 4 | 160.9 (9.3-2793.8) | 4.8561169 | 0.0000118 |
| (MDC-STM-002-0.6) | Control | 126 | 4 | 29.5 (1.6-528.9) | 3.1984047 | 0.0120573 |
| (MDC-STM-003-0.6) | Control | 126 | 4 | 156.2 (9-2712.6) | 4.8271523 | 0.0000137 |
| (MDC-STM-003-1.5) | Control | 126 | 4 | 215.2 (12.4-3727.6) | 5.1371852 | 0.0000028 |
| (MDC-STM-001-0.6) | Control | BI | 10 | 0.6 (0.2-2.3) | -0.9559426 | 0.8747113 |
| (MDC-STM-002-0.6) | Control | BI | 10 | 0.5 (0.1-1.8) | -1.5706514 | 0.5163317 |
| (MDC-STM-003-0.6) | Control | BI | 10 | 0.5 (0.2-1.9) | -1.3559929 | 0.6559403 |
| (MDC-STM-003-1.5) | Control | BI | 10 | 0.4 (0.1-1.5) | -1.9615231 | 0.2850212 |
| (MDC-STM-001-0.6) | Control | 2 | 10 | 22.3 (8.1-61.5) | 8.3298780 | 0.0000000 |
| (MDC-STM-002-0.6) | Control | 2 | 10 | 44.4 (16.1-122.5) | 10.2014974 | 0.0000000 |
| (MDC-STM-003-0.6) | Control | 2 | 10 | 44.1 (16-121.6) | 10.1883342 | 0.0000000 |
| (MDC-STM-003-1.5) | Control | 2 | 10 | 40.8 (14.8-112.8) | 9.9635258 | 0.0000000 |
| (MDC-STM-001-0.6) | Control | 17 | 10 | 22.1 (8.7-56.1) | 9.0876376 | 0.0000000 |
| (MDC-STM-002-0.6) | Control | 17 | 10 | 19.9 (7.9-50.5) | 8.7831843 | 0.0000000 |
| (MDC-STM-003-0.6) | Control | 17 | 10 | 19.5 (7.7-49.5) | 8.7173277 | 0.0000000 |
| (MDC-STM-003-1.5) | Control | 17 | 10 | 22.9 (9-57.9) | 9.1788002 | 0.0000000 |
| (MDC-STM-001-0.6) | Control | 21 | 10 | 19.4 (7.7-48.6) | 8.8005338 | 0.0000000 |
| (MDC-STM-002-0.6) | Control | 21 | 10 | 16.7 (6.7-42) | 8.3551000 | 0.0000000 |
| (MDC-STM-003-0.6) | Control | 21 | 10 | 20.7 (8.3-52) | 8.9892405 | 0.0000000 |
| (MDC-STM-003-1.5) | Control | 21 | 10 | 21.3 (8.5-53.5) | 9.0828836 | 0.0000000 |
| (MDC-STM-001-0.6) | Control | 42 | 10 | 12.1 (4.6-31.9) | 7.0314926 | 0.0000000 |
| (MDC-STM-002-0.6) | Control | 42 | 10 | 7 (2.6-18.4) | 5.4504723 | 0.0000005 |
| (MDC-STM-003-0.6) | Control | 42 | 10 | 8.9 (3.4-23.5) | 6.1803507 | 0.0000000 |
| (MDC-STM-003-1.5) | Control | 42 | 10 | 12.3 (4.7-32.5) | 7.0804487 | 0.0000000 |
| (MDC-STM-001-0.6) | Control | 56 | 10 | 14.4 (5.6-37.2) | 7.6531475 | 0.0000000 |
| (MDC-STM-002-0.6) | Control | 56 | 10 | 7.2 (2.8-18.7) | 5.6511352 | 0.0000002 |
| (MDC-STM-003-0.6) | Control | 56 | 10 | 14.5 (5.6-37.7) | 7.6599087 | 0.0000000 |
| (MDC-STM-003-1.5) | Control | 56 | 10 | 21.2 (8.2-54.9) | 8.7464041 | 0.0000000 |
| (MDC-STM-001-0.6) | Control | 70 | 10 | 12.3 (4.8-31.6) | 7.2559840 | 0.0000000 |
| (MDC-STM-002-0.6) | Control | 70 | 10 | 3.2 (1.2-8.5) | 3.2928640 | 0.0087904 |
| (MDC-STM-003-0.6) | Control | 70 | 10 | 9 (3.5-23.2) | 6.3074379 | 0.0000000 |
| (MDC-STM-003-1.5) | Control | 70 | 10 | 15.7 (6.1-40.3) | 7.9399520 | 0.0000000 |
| (MDC-STM-001-0.6) | Control | 84 | 10 | 8 (3.2-20.3) | 6.1630229 | 0.0000000 |
| (MDC-STM-002-0.6) | Control | 84 | 10 | 2.4 (1-6.3) | 2.5834060 | 0.0734027 |
| (MDC-STM-003-0.6) | Control | 84 | 10 | 5.7 (2.2-14.4) | 5.0764744 | 0.0000038 |
| (MDC-STM-003-1.5) | Control | 84 | 10 | 10.1 (4-25.4) | 6.8093812 | 0.0000000 |
| (MDC-STM-001-0.6) | Control | 98 | 10 | 10.1 (3.9-26.2) | 6.5853039 | 0.0000000 |
| (MDC-STM-002-0.6) | Control | 98 | 10 | 2.1 (0.8-5.5) | 1.9907963 | 0.2703929 |
| (MDC-STM-003-0.6) | Control | 98 | 10 | 5.7 (2.2-15.1) | 4.9284663 | 0.0000082 |
| (MDC-STM-003-1.5) | Control | 98 | 10 | 9.6 (3.7-25) | 6.4097776 | 0.0000000 |
| (MDC-STM-001-0.6) | Control | 112 | 10 | 10.9 (4-29.9) | 6.4398595 | 0.0000000 |
| (MDC-STM-002-0.6) | Control | 112 | 10 | 3.3 (1.2-9.3) | 3.1723213 | 0.0131335 |
| (MDC-STM-003-0.6) | Control | 112 | 10 | 10.7 (3.9-29.5) | 6.3876542 | 0.0000000 |
| (MDC-STM-003-1.5) | Control | 112 | 10 | 11.5 (4.2-31.6) | 6.5809723 | 0.0000000 |
| (MDC-STM-001-0.6) | Control | 126 | 10 | 34.9 (9.3-131.3) | 7.3172285 | 0.0000000 |
| (MDC-STM-002-0.6) | Control | 126 | 10 | 10.5 (2.7-40.3) | 4.7708941 | 0.0000181 |
| (MDC-STM-003-0.6) | Control | 126 | 10 | 33.2 (8.8-125.1) | 7.2074558 | 0.0000000 |
| (MDC-STM-003-1.5) | Control | 126 | 10 | 45.5 (12.1-170.7) | 7.8755092 | 0.0000000 |
| (MDC-STM-001-0.6) | Control | BI | 30 | 1.1 (0.5-2.3) | 0.3311432 | 0.9974111 |
| (MDC-STM-002-0.6) | Control | BI | 30 | 1 (0.5-2.1) | -0.0125675 | 1.0000000 |
| (MDC-STM-003-0.6) | Control | BI | 30 | 0.8 (0.4-1.8) | -0.6516467 | 0.9663867 |
| (MDC-STM-003-1.5) | Control | BI | 30 | 0.9 (0.4-1.9) | -0.4079445 | 0.9941915 |
| (MDC-STM-001-0.6) | Control | 2 | 30 | 7.4 (3.5-15.7) | 7.2400202 | 0.0000000 |
| (MDC-STM-002-0.6) | Control | 2 | 30 | 18.3 (8.6-38.9) | 10.5353421 | 0.0000000 |
| (MDC-STM-003-0.6) | Control | 2 | 30 | 18.4 (8.7-39.1) | 10.5658195 | 0.0000000 |
| (MDC-STM-003-1.5) | Control | 2 | 30 | 12.6 (5.9-26.8) | 9.1477425 | 0.0000000 |
| (MDC-STM-001-0.6) | Control | 17 | 30 | 10.6 (5.1-22.3) | 8.7066586 | 0.0000000 |
| (MDC-STM-002-0.6) | Control | 17 | 30 | 9.4 (4.5-19.7) | 8.2714953 | 0.0000000 |
| (MDC-STM-003-0.6) | Control | 17 | 30 | 9.4 (4.5-19.6) | 8.2404428 | 0.0000000 |
| (MDC-STM-003-1.5) | Control | 17 | 30 | 11.8 (5.6-24.7) | 9.0870759 | 0.0000000 |
| (MDC-STM-001-0.6) | Control | 21 | 30 | 10.1 (4.8-21.2) | 8.5489587 | 0.0000000 |
| (MDC-STM-002-0.6) | Control | 21 | 30 | 7.4 (3.5-15.4) | 7.3709439 | 0.0000000 |
| (MDC-STM-003-0.6) | Control | 21 | 30 | 10.7 (5.1-22.4) | 8.7404073 | 0.0000000 |
| (MDC-STM-003-1.5) | Control | 21 | 30 | 11.6 (5.6-24.3) | 9.0550481 | 0.0000000 |
| (MDC-STM-001-0.6) | Control | 42 | 30 | 5.9 (2.7-12.7) | 6.3153947 | 0.0000000 |
| (MDC-STM-002-0.6) | Control | 42 | 30 | 3.8 (1.8-8.2) | 4.7956175 | 0.0000160 |
| (MDC-STM-003-0.6) | Control | 42 | 30 | 4.7 (2.2-10.2) | 5.5638129 | 0.0000003 |
| (MDC-STM-003-1.5) | Control | 42 | 30 | 6.8 (3.2-14.6) | 6.8245889 | 0.0000000 |
| (MDC-STM-001-0.6) | Control | 56 | 30 | 5.5 (2.6-11.7) | 6.2391368 | 0.0000000 |
| (MDC-STM-002-0.6) | Control | 56 | 30 | 3.1 (1.5-6.6) | 4.1792820 | 0.0002827 |
| (MDC-STM-003-0.6) | Control | 56 | 30 | 6.8 (3.2-14.3) | 6.9961241 | 0.0000000 |
| (MDC-STM-003-1.5) | Control | 56 | 30 | 7.1 (3.4-15.1) | 7.1113294 | 0.0000000 |
| (MDC-STM-001-0.6) | Control | 70 | 30 | 3.5 (1.7-7.2) | 4.6389669 | 0.0000345 |
| (MDC-STM-002-0.6) | Control | 70 | 30 | 1.4 (0.7-2.9) | 1.2618253 | 0.7146990 |
| (MDC-STM-003-0.6) | Control | 70 | 30 | 2.7 (1.3-5.7) | 3.7237241 | 0.0018361 |
| (MDC-STM-003-1.5) | Control | 70 | 30 | 4.8 (2.3-9.9) | 5.8158608 | 0.0000001 |
| (MDC-STM-001-0.6) | Control | 84 | 30 | 3.6 (1.7-7.6) | 4.8051976 | 0.0000153 |
| (MDC-STM-002-0.6) | Control | 84 | 30 | 1.5 (0.7-3.2) | 1.5836401 | 0.5079043 |
| (MDC-STM-003-0.6) | Control | 84 | 30 | 2.5 (1.2-5.1) | 3.3574350 | 0.0070421 |
| (MDC-STM-003-1.5) | Control | 84 | 30 | 4.2 (2-8.7) | 5.3227185 | 0.0000010 |
| (MDC-STM-001-0.6) | Control | 98 | 30 | 3.8 (1.8-8.1) | 4.8907639 | 0.0000099 |
| (MDC-STM-002-0.6) | Control | 98 | 30 | 1.4 (0.6-2.9) | 1.1222767 | 0.7948357 |
| (MDC-STM-003-0.6) | Control | 98 | 30 | 2.8 (1.3-6) | 3.7662399 | 0.0015562 |
| (MDC-STM-003-1.5) | Control | 98 | 30 | 4.4 (2.1-9.3) | 5.3605195 | 0.0000008 |
| (MDC-STM-001-0.6) | Control | 112 | 30 | 3.7 (1.8-7.8) | 4.7881568 | 0.0000166 |
| (MDC-STM-002-0.6) | Control | 112 | 30 | 1.5 (0.7-3.2) | 1.4979596 | 0.5637848 |
| (MDC-STM-003-0.6) | Control | 112 | 30 | 3.7 (1.8-7.9) | 4.8328687 | 0.0000133 |
| (MDC-STM-003-1.5) | Control | 112 | 30 | 4.4 (2.1-9.3) | 5.4279176 | 0.0000006 |
| (MDC-STM-001-0.6) | Control | 126 | 30 | 3.9 (1.9-8.3) | 5.0130665 | 0.0000053 |
| (MDC-STM-002-0.6) | Control | 126 | 30 | 1.7 (0.8-3.6) | 1.8970277 | 0.3188277 |
| (MDC-STM-003-0.6) | Control | 126 | 30 | 4.1 (2-8.7) | 5.1720047 | 0.0000023 |
| (MDC-STM-003-1.5) | Control | 126 | 30 | 4.6 (2.2-9.7) | 5.5941222 | 0.0000002 |

B.
| Formulation | Treatment arm | DAI | Follow-up<br>(days) | HR | CI95% | z.ratio | p.value |
| --- | --- | --- | --- | --- | --- | --- | --- |
| mdc-STM-001(0.6) | Control | BI | 4 | 0.4 | (0.1-2.3) | -1.4525092 | 0.5935112 |
| mdc-STM-002(0.6) | Control | BI | 4 | 0.3 | (0-1.9) | -1.7950230 | 0.3763965 |
| mdc-STM-003(0.6) | Control | BI | 4 | 0.6 | (0.1-2.6) | -0.9988111 | 0.8560128 |
| mdc-STM-003(1.5) | Control | BI | 4 | 0.5 | (0.1-2.7) | -1.0328193 | 0.8402120 |
| mdc-STM-001(0.6) | Control | 2 | 4 | 75.1 | (22.9-246) | 9.9245770 | 0.0000000 |
| mdc-STM-002(0.6) | Control | 2 | 4 | 72.9 | (22.3-239) | 9.8587828 | 0.0000000 |
| mdc-STM-003(0.6) | Control | 2 | 4 | 101.1 | (30.9-331.1) | 10.6117691 | 0.0000000 |
| mdc-STM-003(1.5) | Control | 2 | 4 | 91 | (27.8-298) | 10.3768245 | 0.0000000 |
| mdc-STM-001(0.6) | Control | 7 | 4 | 134.2 | (28.1-640.2) | 8.5530652 | 0.0000000 |
| mdc-STM-002(0.6) | Control | 7 | 4 | 169.1 | (35.5-806.5) | 8.9586219 | 0.0000000 |
| mdc-STM-003(0.6) | Control | 7 | 4 | 178.3 | (37.4-850.7) | 9.0478567 | 0.0000000 |
| mdc-STM-003(1.5) | Control | 7 | 4 | 255.2 | (53.5-1217) | 9.6791985 | 0.0000000 |
| mdc-STM-001(0.6) | Control | 17 | 4 | 53.1 | (15.5-181.7) | 8.8085244 | 0.0000000 |
| mdc-STM-002(0.6) | Control | 17 | 4 | 35.5 | (10.4-121.7) | 7.9027325 | 0.0000000 |
| mdc-STM-003(0.6) | Control | 17 | 4 | 50.7 | (14.8-173.6) | 8.7020934 | 0.0000000 |
| mdc-STM-003(1.5) | Control | 17 | 4 | 69.4 | (20.3-237.1) | 9.4072719 | 0.0000000 |
| mdc-STM-001(0.6) | Control | 21 | 4 | 62.4 | (17.3-225.1) | 8.7889633 | 0.0000000 |
| mdc-STM-002(0.6) | Control | 21 | 4 | 41.4 | (11.4-149.7) | 7.8924001 | 0.0000000 |
| mdc-STM-003(0.6) | Control | 21 | 4 | 45.2 | (12.5-163.7) | 8.0827196 | 0.0000000 |
| mdc-STM-003(1.5) | Control | 21 | 4 | 73.1 | (20.3-263.7) | 9.1269221 | 0.0000000 |
| mdc-STM-001(0.6) | Control | 28 | 4 | 97 | (20.3-463.2) | 7.9817790 | 0.0000000 |
| mdc-STM-002(0.6) | Control | 28 | 4 | 50.1 | (10.5-240.3) | 6.8120194 | 0.0000000 |
| mdc-STM-003(0.6) | Control | 28 | 4 | 51.5 | (10.7-246.8) | 6.8614600 | 0.0000000 |
| mdc-STM-003(1.5) | Control | 28 | 4 | 77 | (16.1-368) | 7.5763840 | 0.0000000 |
| mdc-STM-001(0.6) | Control | 42 | 4 | 25.3 | (8.2-77.9) | 7.8344226 | 0.0000000 |
| mdc-STM-002(0.6) | Control | 42 | 4 | 8.2 | (2.6-25.8) | 4.9966923 | 0.0000058 |
| mdc-STM-003(0.6) | Control | 42 | 4 | 16.7 | (5.4-51.8) | 6.7820428 | 0.0000000 |
| mdc-STM-003(1.5) | Control | 42 | 4 | 28.4 | (9.2-87.5) | 8.1068411 | 0.0000000 |
| mdc-STM-001(0.6) | Control | 56 | 4 | 78.8 | (13.6-454.9) | 6.7925446 | 0.0000000 |
| mdc-STM-002(0.6) | Control | 56 | 4 | 20.4 | (3.4-121) | 4.6202949 | 0.0000377 |
| mdc-STM-003(0.6) | Control | 56 | 4 | 55.9 | (9.6-324.8) | 6.2355142 | 0.0000000 |
| mdc-STM-003(1.5) | Control | 56 | 4 | 115.4 | (20-665.7) | 7.3936684 | 0.0000000 |
| mdc-STM-001(0.6) | Control | 70 | 4 | 14.3 | (4.1-49.8) | 5.8249608 | 0.0000001 |
| mdc-STM-002(0.6) | Control | 70 | 4 | 4.2 | (1.1-15.4) | 3.0047931 | 0.0223285 |
| mdc-STM-003(0.6) | Control | 70 | 4 | 14.7 | (4.2-51.1) | 5.8808958 | 0.0000000 |
| mdc-STM-003(1.5) | Control | 70 | 4 | 26.8 | (7.8-92.5) | 7.2458816 | 0.0000000 |
| mdc-STM-001(0.6) | Control | 84 | 4 | 8.8 | (2.6-29.8) | 4.8796195 | 0.0000105 |
| mdc-STM-002(0.6) | Control | 84 | 4 | 1.4 | (0.4-5.7) | 0.7288331 | 0.9498655 |
| mdc-STM-003(0.6) | Control | 84 | 4 | 13.7 | (4.1-45.9) | 5.9298691 | 0.0000000 |
| mdc-STM-003(1.5) | Control | 84 | 4 | 16.9 | (5.1-56.1) | 6.4215395 | 0.0000000 |
| mdc-STM-001(0.6) | Control | 91 | 4 | 43.8 | (5.4-353.9) | 4.9315684 | 0.0000081 |
| mdc-STM-002(0.6) | Control | 91 | 4 | 8.3 | (0.9-73.3) | 2.6608551 | 0.0599267 |
| mdc-STM-003(0.6) | Control | 91 | 4 | 52.4 | (6.5-422.1) | 5.1774268 | 0.0000022 |
| mdc-STM-003(1.5) | Control | 91 | 4 | 68.5 | (8.5-550.5) | 5.5334036 | 0.0000003 |
| mdc-STM-001(0.6) | Control | 98 | 4 | 8.7 | (2.3-32.5) | 4.4604133 | 0.0000801 |
| mdc-STM-002(0.6) | Control | 98 | 4 | 2.1 | (0.5-8.9) | 1.3968548 | 0.6296925 |
| mdc-STM-003(0.6) | Control | 98 | 4 | 8.5 | (2.3-31.8) | 4.4446592 | 0.0000861 |

| Formulation | Treatment arm | DAI | Follow-up<br>(days) | HR CI95% | z.ratio | p.value |
| --- | --- | --- | --- | --- | --- | --- |
| mdc-STM-003(1.5) | Control | 98 | 4 | 14.8 (4-54.2) | 5.6493493 | 0.0000002 |
| mdc-STM-001(0.6) | Control | 105 | 4 | 7.1 (2.2-23.3) | 4.5143215 | 0.0000623 |
| mdc-STM-002(0.6) | Control | 105 | 4 | 1.3 (0.3-4.9) | 0.4836030 | 0.9888789 |
| mdc-STM-003(0.6) | Control | 105 | 4 | 6.8 (2.1-22.2) | 4.4256915 | 0.0000940 |
| mdc-STM-003(1.5) | Control | 105 | 4 | 14.8 (4.6-47.2) | 6.3098761 | 0.0000000 |
| mdc-STM-001(0.6) | Control | 112 | 4 | 5.6 (1.6-20.3) | 3.6722060 | 0.0022378 |
| mdc-STM-002(0.6) | Control | 112 | 4 | 2.2 (0.6-8.8) | 1.5855045 | 0.5066966 |
| mdc-STM-003(0.6) | Control | 112 | 4 | 7.1 (2-25.1) | 4.2230989 | 0.0002335 |
| mdc-STM-003(1.5) | Control | 112 | 4 | 9.4 (2.7-33) | 4.8654735 | 0.0000113 |
| mdc-STM-001(0.6) | Control | 119 | 4 | 15.5 (3.2-76.1) | 4.6954432 | 0.0000262 |
| mdc-STM-002(0.6) | Control | 119 | 4 | 3.4 (0.6-18.7) | 1.9371514 | 0.2975453 |
| mdc-STM-003(0.6) | Control | 119 | 4 | 18.3 (3.8-89.3) | 5.0046730 | 0.0000055 |
| mdc-STM-003(1.5) | Control | 119 | 4 | 24.9 (5.1-120.6) | 5.5529337 | 0.0000003 |
| mdc-STM-001(0.6) | Control | 126 | 4 | 42.1 (5.2-340.3) | 4.8779078 | 0.0000106 |
| mdc-STM-002(0.6) | Control | 126 | 4 | 5.5 (0.6-51.2) | 2.1031035 | 0.2186133 |
| mdc-STM-003(0.6) | Control | 126 | 4 | 36.3 (4.5-293.7) | 4.6872015 | 0.0000273 |
| mdc-STM-003(1.5) | Control | 126 | 4 | 56.5 (7-454.2) | 5.2765308 | 0.0000013 |
| mdc-STM-001(0.6) | Control | BI | 10 | 0.5 (0.2-1.4) | -1.8412832 | 0.3497023 |
| mdc-STM-002(0.6) | Control | BI | 10 | 0.6 (0.2-1.6) | -1.4389514 | 0.6023572 |
| mdc-STM-003(0.6) | Control | BI | 10 | 0.7 (0.3-1.8) | -1.1321622 | 0.7895059 |
| mdc-STM-003(1.5) | Control | BI | 10 | 0.7 (0.3-1.9) | -0.9502176 | 0.8771030 |
| mdc-STM-001(0.6) | Control | 2 | 10 | 23.7 (10.5-53.7) | 10.5671244 | 0.0000000 |
| mdc-STM-002(0.6) | Control | 2 | 10 | 24.5 (10.8-55.4) | 10.6674750 | 0.0000000 |
| mdc-STM-003(0.6) | Control | 2 | 10 | 34.7 (15.3-78.6) | 11.8402697 | 0.0000000 |
| mdc-STM-003(1.5) | Control | 2 | 10 | 31.4 (13.9-71.1) | 11.5091218 | 0.0000000 |
| mdc-STM-001(0.6) | Control | 7 | 10 | 21.8 (9.5-49.9) | 10.1539733 | 0.0000000 |
| mdc-STM-002(0.6) | Control | 7 | 10 | 28.2 (12.3-64.4) | 10.9963707 | 0.0000000 |
| mdc-STM-003(0.6) | Control | 7 | 10 | 29.5 (12.9-67.5) | 11.1369242 | 0.0000000 |
| mdc-STM-003(1.5) | Control | 7 | 10 | 43.3 (18.9-99) | 12.4087000 | 0.0000000 |
| mdc-STM-001(0.6) | Control | 17 | 10 | 12.4 (5.6-27.6) | 8.6218624 | 0.0000000 |
| mdc-STM-002(0.6) | Control | 17 | 10 | 8.3 (3.7-18.5) | 7.2272830 | 0.0000000 |
| mdc-STM-003(0.6) | Control | 17 | 10 | 12.4 (5.6-27.5) | 8.6164337 | 0.0000000 |
| mdc-STM-003(1.5) | Control | 17 | 10 | 17.5 (7.9-38.8) | 9.7930817 | 0.0000000 |
| mdc-STM-001(0.6) | Control | 21 | 10 | 11.1 (5.1-24.4) | 8.3818904 | 0.0000000 |
| mdc-STM-002(0.6) | Control | 21 | 10 | 8 (3.6-17.6) | 7.2161276 | 0.0000000 |
| mdc-STM-003(0.6) | Control | 21 | 10 | 9.1 (4.1-19.9) | 7.6585295 | 0.0000000 |
| mdc-STM-003(1.5) | Control | 21 | 10 | 12.9 (5.9-28.4) | 8.8957932 | 0.0000000 |
| mdc-STM-001(0.6) | Control | 28 | 10 | 19.3 (8.1-45.7) | 9.3607437 | 0.0000000 |
| mdc-STM-002(0.6) | Control | 28 | 10 | 10.5 (4.4-25.1) | 7.4189722 | 0.0000000 |
| mdc-STM-003(0.6) | Control | 28 | 10 | 10.6 (4.5-25.3) | 7.4419849 | 0.0000000 |
| mdc-STM-003(1.5) | Control | 28 | 10 | 15.9 (6.7-37.6) | 8.7267934 | 0.0000000 |
| mdc-STM-001(0.6) | Control | 42 | 10 | 10.4 (4.5-24.1) | 7.6317937 | 0.0000000 |
| mdc-STM-002(0.6) | Control | 42 | 10 | 3.8 (1.6-8.9) | 4.2488554 | 0.0002084 |
| mdc-STM-003(0.6) | Control | 42 | 10 | 7.2 (3.1-16.8) | 6.3764610 | 0.0000000 |
| mdc-STM-003(1.5) | Control | 42 | 10 | 12.9 (5.6-29.7) | 8.3139612 | 0.0000000 |
| mdc-STM-001(0.6) | Control | 56 | 10 | 28.4 (10.1-80.2) | 8.8128648 | 0.0000000 |
| mdc-STM-002(0.6) | Control | 56 | 10 | 8.2 (2.8-23.6) | 5.4098301 | 0.0000006 |
| mdc-STM-003(0.6) | Control | 56 | 10 | 19.1 (6.7-54.4) | 7.7040423 | 0.0000000 |
| mdc-STM-003(1.5) | Control | 56 | 10 | 39.7 (14.1-111.8) | 9.6938839 | 0.0000000 |
| mdc-STM-001(0.6) | Control | 70 | 10 | 12.9 (4.6-36) | 6.8149255 | 0.0000000 |
| mdc-STM-002(0.6) | Control | 70 | 10 | 4.4 (1.5-12.6) | 3.8067849 | 0.0013268 |
| mdc-STM-003(0.6) | Control | 70 | 10 | 12.5 (4.5-34.9) | 6.7040907 | 0.0000000 |
| mdc-STM-003(1.5) | Control | 70 | 10 | 21.4 (7.7-59.5) | 8.1851871 | 0.0000000 |
| mdc-STM-001(0.6) | Control | 84 | 10 | 2.9 (1.3-6.7) | 3.5562395 | 0.0034569 |
| mdc-STM-002(0.6) | Control | 84 | 10 | 1.2 (0.5-2.9) | 0.6589596 | 0.9650042 |
| mdc-STM-003(0.6) | Control | 84 | 10 | 4.2 (1.9-9.6) | 4.8024106 | 0.0000155 |
| mdc-STM-003(1.5) | Control | 84 | 10 | 5.3 (2.4-12.1) | 5.6207999 | 0.0000002 |
| mdc-STM-001(0.6) | Control | 91 | 10 | 5.7 (2.3-14) | 5.3194529 | 0.0000010 |
| mdc-STM-002(0.6) | Control | 91 | 10 | 2.1 (0.8-5.3) | 2.1260924 | 0.2088806 |
| mdc-STM-003(0.6) | Control | 91 | 10 | 6.6 (2.7-16.2) | 5.7875928 | 0.0000001 |
| mdc-STM-003(1.5) | Control | 91 | 10 | 7.7 (3.2-18.8) | 6.2545248 | 0.0000000 |
| mdc-STM-001(0.6) | Control | 98 | 10 | 5.4 (2.1-13.8) | 4.9224411 | 0.0000085 |
| mdc-STM-002(0.6) | Control | 98 | 10 | 2.4 (0.9-6.3) | 2.4565738 | 0.1007138 |
| mdc-STM-003(0.6) | Control | 98 | 10 | 5.4 (2.1-13.7) | 4.9189045 | 0.0000086 |
| mdc-STM-003(1.5) | Control | 98 | 10 | 8.1 (3.2-20.3) | 6.1510916 | 0.0000000 |
| mdc-STM-001(0.6) | Control | 105 | 10 | 3.8 (1.6-9.1) | 4.1562642 | 0.0003123 |
| mdc-STM-002(0.6) | Control | 105 | 10 | 1.8 (0.7-4.3) | 1.7014145 | 0.4329995 |
| mdc-STM-003(0.6) | Control | 105 | 10 | 4.2 (1.8-10.1) | 4.5216255 | 0.0000602 |
| mdc-STM-003(1.5) | Control | 105 | 10 | 9.2 (3.9-21.7) | 7.0669937 | 0.0000000 |
| mdc-STM-001(0.6) | Control | 112 | 10 | 3.6 (1.5-8.7) | 3.8574151 | 0.0010845 |
| mdc-STM-002(0.6) | Control | 112 | 10 | 2.1 (0.8-5.2) | 2.1245354 | 0.2095304 |
| mdc-STM-003(0.6) | Control | 112 | 10 | 4.2 (1.7-10.1) | 4.3546434 | 0.0001299 |
| mdc-STM-003(1.5) | Control | 112 | 10 | 4.4 (1.8-10.8) | 4.5611161 | 0.0000500 |
| mdc-STM-001(0.6) | Control | 119 | 10 | 6.2 (2.3-16.6) | 5.0086104 | 0.0000054 |
| mdc-STM-002(0.6) | Control | 119 | 10 | 2.2 (0.8-6.2) | 2.0603323 | 0.2375089 |
| mdc-STM-003(0.6) | Control | 119 | 10 | 6.2 (2.3-16.7) | 5.0217955 | 0.0000051 |
| mdc-STM-003(1.5) | Control | 119 | 10 | 9.9 (3.7-26.5) | 6.4010372 | 0.0000000 |
| mdc-STM-001(0.6) | Control | 126 | 10 | 8.6 (3.3-22) | 6.2106643 | 0.0000000 |
| mdc-STM-002(0.6) | Control | 126 | 10 | 2.2 (0.8-6.1) | 2.1991847 | 0.1799034 |
| mdc-STM-003(0.6) | Control | 126 | 10 | 6.9 (2.7-17.7) | 5.5486393 | 0.0000003 |
| mdc-STM-003(1.5) | Control | 126 | 10 | 9.4 (3.6-24) | 6.4624035 | 0.0000000 |
| mdc-STM-001(0.6) | Control | BI | 30 | 0.9 (0.5-1.7) | -0.4413259 | 0.9921471 |
| mdc-STM-002(0.6) | Control | BI | 30 | 0.8 (0.4-1.6) | -0.7137346 | 0.9534396 |
| mdc-STM-003(0.6) | Control | BI | 30 | 0.7 (0.4-1.4) | -1.3516314 | 0.6587202 |
| mdc-STM-003(1.5) | Control | BI | 30 | 0.7 (0.4-1.4) | -1.3346408 | 0.6695032 |
| mdc-STM-001(0.6) | Control | 2 | 30 | 9.5 (5-18.1) | 9.5963295 | 0.0000000 |
| mdc-STM-002(0.6) | Control | 2 | 30 | 12.6 (6.7-24) | 10.7821368 | 0.0000000 |
| mdc-STM-003(0.6) | Control | 2 | 30 | 15.2 (8-28.9) | 11.5763541 | 0.0000000 |
| mdc-STM-003(1.5) | Control | 2 | 30 | 15.5 (8.2-29.5) | 11.6599028 | 0.0000000 |
| mdc-STM-001(0.6) | Control | 7 | 30 | 9.9 (5.2-18.9) | 9.7065456 | 0.0000000 |
| mdc-STM-002(0.6) | Control | 7 | 30 | 11.7 (6.1-22.2) | 10.3959345 | 0.0000000 |
| mdc-STM-003(0.6) | Control | 7 | 30 | 13.1 (6.9-24.9) | 10.8754247 | 0.0000000 |
| mdc-STM-003(1.5) | Control | 7 | 30 | 20.5 (10.7-39) | 12.7590423 | 0.0000000 |
| mdc-STM-001(0.6) | Control | 17 | 30 | 7.1 (3.8-13.5) | 8.3577625 | 0.0000000 |
| mdc-STM-002(0.6) | Control | 17 | 30 | 4.3 (2.3-8.2) | 6.2506357 | 0.0000000 |
| mdc-STM-003(0.6) | Control | 17 | 30 | 6.5 (3.4-12.4) | 7.9786739 | 0.0000000 |
| mdc-STM-003(1.5) | Control | 17 | 30 | 10.6 (5.6-20.2) | 10.0472269 | 0.0000000 |
| mdc-STM-001(0.6) | Control | 21 | 30 | 7.6 (4-14.4) | 8.6089502 | 0.0000000 |
| mdc-STM-002(0.6) | Control | 21 | 30 | 5 (2.6-9.5) | 6.8561647 | 0.0000000 |
| mdc-STM-003(0.6) | Control | 21 | 30 | 5.2 (2.7-9.9) | 6.9905957 | 0.0000000 |
| mdc-STM-003(1.5) | Control | 21 | 30 | 8.9 (4.7-16.9) | 9.2674039 | 0.0000000 |
| mdc-STM-001(0.6) | Control | 28 | 30 | 6.8 (3.6-13) | 8.1629184 | 0.0000000 |
| mdc-STM-002(0.6) | Control | 28 | 30 | 3.5 (1.8-6.7) | 5.3356150 | 0.0000009 |
| mdc-STM-003(0.6) | Control | 28 | 30 | 3.5 (1.9-6.7) | 5.3392816 | 0.0000009 |
| mdc-STM-003(1.5) | Control | 28 | 30 | 5.6 (2.9-10.6) | 7.2901991 | 0.0000000 |
| mdc-STM-001(0.6) | Control | 42 | 30 | 5.5 (2.9-10.7) | 7.0734936 | 0.0000000 |
| mdc-STM-002(0.6) | Control | 42 | 30 | 2.2 (1.1-4.2) | 3.1845824 | 0.0126174 |
| mdc-STM-003(0.6) | Control | 42 | 30 | 4.4 (2.3-8.4) | 6.0755004 | 0.0000000 |
| mdc-STM-003(1.5) | Control | 42 | 30 | 6.3 (3.2-12.1) | 7.5618084 | 0.0000000 |
| mdc-STM-001(0.6) | Control | 56 | 30 | 4.8 (2.5-9.1) | 6.6095855 | 0.0000000 |
| mdc-STM-002(0.6) | Control | 56 | 30 | 1.8 (0.9-3.4) | 2.4143863 | 0.1113955 |
| mdc-STM-003(0.6) | Control | 56 | 30 | 3.4 (1.8-6.5) | 5.1304999 | 0.0000029 |
| mdc-STM-003(1.5) | Control | 56 | 30 | 7 (3.7-13.4) | 8.2612435 | 0.0000000 |
| mdc-STM-001(0.6) | Control | 70 | 30 | 2.8 (1.5-5.3) | 4.3426810 | 0.0001371 |
| mdc-STM-002(0.6) | Control | 70 | 30 | 1.5 (0.8-2.8) | 1.5709813 | 0.5161174 |
| mdc-STM-003(0.6) | Control | 70 | 30 | 2.5 (1.3-4.7) | 3.8103196 | 0.0013084 |
| mdc-STM-003(1.5) | Control | 70 | 30 | 4 (2.1-7.6) | 5.8398598 | 0.0000001 |
| mdc-STM-001(0.6) | Control | 84 | 30 | 1.6 (0.9-3.1) | 2.1039597 | 0.2182455 |
| mdc-STM-002(0.6) | Control | 84 | 30 | 1.1 (0.6-2.2) | 0.5889268 | 0.9767710 |
| mdc-STM-003(0.6) | Control | 84 | 30 | 2.4 (1.3-4.6) | 3.8137967 | 0.0012905 |
| mdc-STM-003(1.5) | Control | 84 | 30 | 2.7 (1.4-5.2) | 4.2572940 | 0.0002008 |
| mdc-STM-001(0.6) | Control | 91 | 30 | 1.9 (1-3.7) | 2.7988980 | 0.0410041 |
| mdc-STM-002(0.6) | Control | 91 | 30 | 1.2 (0.6-2.2) | 0.6465748 | 0.9673238 |
| mdc-STM-003(0.6) | Control | 91 | 30 | 2.3 (1.2-4.4) | 3.5590342 | 0.0034214 |
| mdc-STM-003(1.5) | Control | 91 | 30 | 2.5 (1.3-4.9) | 3.9686477 | 0.0006898 |
| mdc-STM-001(0.6) | Control | 98 | 30 | 2 (1.1-3.9) | 3.0181363 | 0.0214296 |
| mdc-STM-002(0.6) | Control | 98 | 30 | 1.2 (0.6-2.3) | 0.8332180 | 0.9203662 |
| mdc-STM-003(0.6) | Control | 98 | 30 | 2 (1.1-3.8) | 2.9714118 | 0.0247237 |
| mdc-STM-003(1.5) | Control | 98 | 30 | 2.5 (1.3-4.8) | 3.9617328 | 0.0007098 |
| mdc-STM-001(0.6) | Control | 105 | 30 | 1.4 (0.8-2.7) | 1.5660574 | 0.5193176 |
| mdc-STM-002(0.6) | Control | 105 | 30 | 1.1 (0.6-2.1) | 0.4346779 | 0.9925898 |
| mdc-STM-003(0.6) | Control | 105 | 30 | 1.7 (0.9-3.3) | 2.3262059 | 0.1365363 |
| mdc-STM-003(1.5) | Control | 105 | 30 | 2.9 (1.5-5.4) | 4.5183891 | 0.0000611 |
| mdc-STM-001(0.6) | Control | 112 | 30 | 1.6 (0.9-3.1) | 2.1173510 | 0.2125465 |
| mdc-STM-002(0.6) | Control | 112 | 30 | 1.3 (0.7-2.4) | 0.9586987 | 0.8735510 |
| mdc-STM-003(0.6) | Control | 112 | 30 | 1.7 (0.9-3.2) | 2.1785290 | 0.1877897 |
| mdc-STM-003(1.5) | Control | 112 | 30 | 2.1 (1.1-4) | 3.1795246 | 0.0128281 |
| mdc-STM-001(0.6) | Control | 119 | 30 | 1.9 (1-3.7) | 2.7986785 | 0.0410296 |
| mdc-STM-002(0.6) | Control | 119 | 30 | 1.3 (0.7-2.5) | 1.1645890 | 0.7716195 |
| mdc-STM-003(0.6) | Control | 119 | 30 | 1.9 (1-3.7) | 2.7812962 | 0.0430913 |
| mdc-STM-003(1.5) | Control | 119 | 30 | 2.6 (1.4-5) | 4.0242898 | 0.0005475 |
| mdc-STM-001(0.6) | Control | 126 | 30 | 2.2 (1.2-4.2) | 3.3472055 | 0.0072961 |
| mdc-STM-002(0.6) | Control | 126 | 30 | 1 (0.5-2) | 0.1792798 | 0.9997694 |
| mdc-STM-003(0.6) | Control | 126 | 30 | 1.9 (1-3.6) | 2.7780905 | 0.0434810 |
| mdc-STM-003(1.5) | Control | 126 | 30 | 2.2 (1.1-4.1) | 3.2770919 | 0.0092731 |

**Supplementary table S9.**
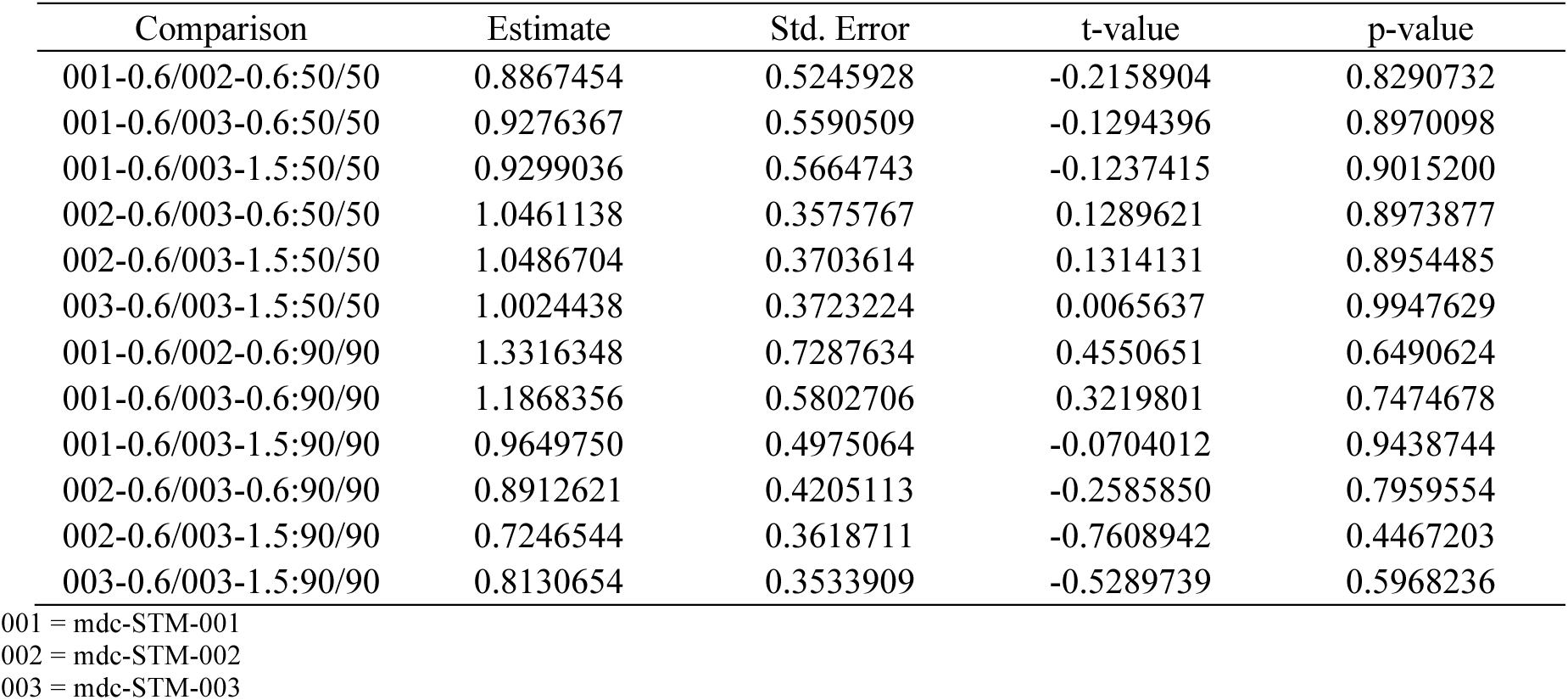
Between formulations comparisons of 4-day LC50 and L90 values for KIS mosquitoes fed on treated cattle.

**Supplementary Table S10.** Between formulations comparisons of 10-day LC50 and LC90 values for KIS mosquitoes fed on treated cattle.

| Comparison | Estimate | Std. Error | t-value | p-value |
| --- | --- | --- | --- | --- |
| 001-0.6/002-0.6:50/50 | 0.8503147 | 0.4901133 | -0.3054095 | 0.7600542 |
| 001-0.6/003-0.6:50/50 | 0.8484447 | 0.4406361 | -0.3439467 | 0.7308864 |
| 001-0.6/003-1.5:50/50 | 0.9622920 | 0.5826786 | -0.0647150 | 0.9484009 |
| 002-0.6/003-0.6:50/50 | 0.9978008 | 0.4234921 | -0.0051931 | 0.9958566 |
| 002-0.6/003-1.5:50/50 | 1.1316892 | 0.5956824 | 0.2210728 | 0.8250358 |
| 003-0.6/003-1.5:50/50 | 1.1341835 | 0.5253544 | 0.2554152 | 0.7984025 |
| 001-0.6/002-0.6:90/90 | 0.9526083 | 0.5708619 | -0.0830179 | 0.9338373 |
| 001-0.6/003-0.6:90/90 | 1.1594824 | 0.5445723 | 0.2928580 | 0.7696307 |
| 001-0.6/003-1.5:90/90 | 1.0292734 | 0.5106321 | 0.0573278 | 0.9542841 |
| 002-0.6/003-0.6:90/90 | 1.2171660 | 0.6993343 | 0.3105324 | 0.7561561 |
| 002-0.6/003-1.5:90/90 | 1.0804792 | 0.6443631 | 0.1248973 | 0.9006048 |
| 003-0.6/003-1.5:90/90 | 0.8877008 | 0.4136421 | -0.2714888 | 0.7860151 |

001 = mdc-STM-001
002 = mdc-STM-002
003 = mdc-STM-003

**Supplementary Table S11.**
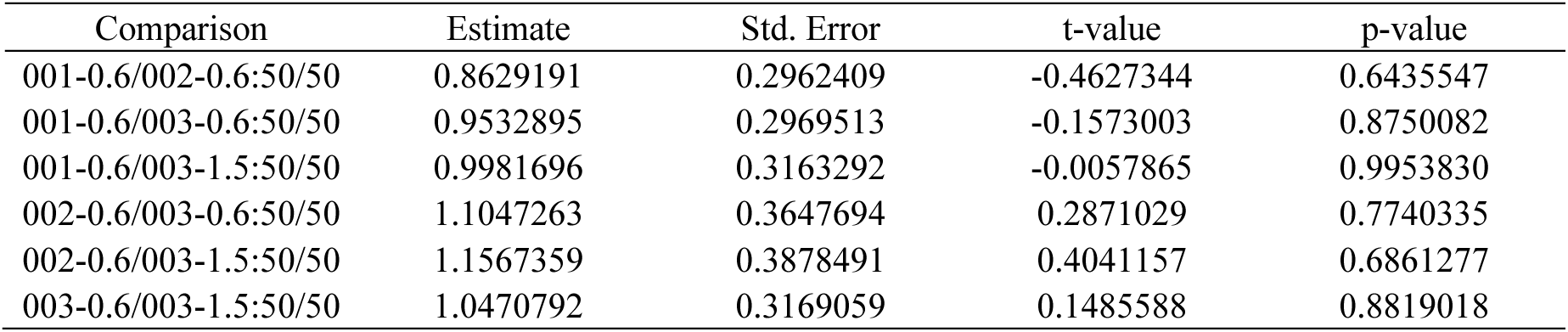
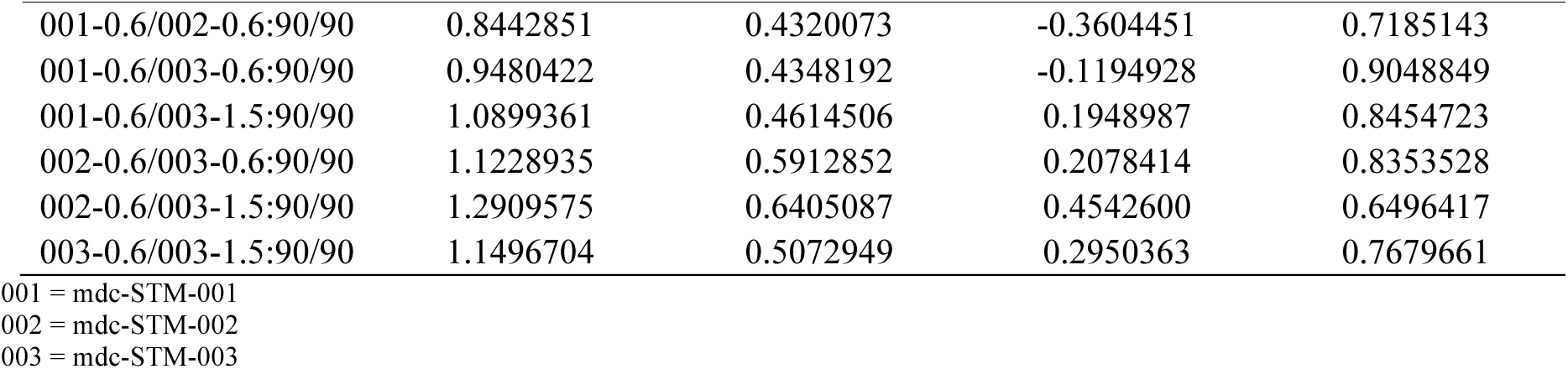
Between formulations comparisons of 4-day LC50 and LC90 values for VK5 mosquitoes fed on treated cattle.

| Comparison | Estimate | Std. Error | t-value | p-value |
| --- | --- | --- | --- | --- |
| 001-0.6/002-0.6:50/50 | 0.8629191 | 0.2962409 | -0.4627344 | 0.6435547 |
| 001-0.6/003-0.6:50/50 | 0.9532895 | 0.2969513 | -0.1573003 | 0.8750082 |
| 001-0.6/003-1.5:50/50 | 0.9981696 | 0.3163292 | -0.0057865 | 0.9953830 |
| 002-0.6/003-0.6:50/50 | 1.1047263 | 0.3647694 | 0.2871029 | 0.7740335 |
| 002-0.6/003-1.5:50/50 | 1.1567359 | 0.3878491 | 0.4041157 | 0.6861277 |
| 003-0.6/003-1.5:50/50 | 1.0470792 | 0.3169059 | 0.1485588 | 0.8819018 |
| 001-0.6/002-0.6:90/90 | 0.8442851 | 0.4320073 | -0.3604451 | 0.7185143 |
| 001-0.6/003-0.6:90/90 | 0.9480422 | 0.4348192 | -0.1194928 | 0.9048849 |
| 001-0.6/003-1.5:90/90 | 1.0899361 | 0.4614506 | 0.1948987 | 0.8454723 |
| 002-0.6/003-0.6:90/90 | 1.1228935 | 0.5912852 | 0.2078414 | 0.8353528 |
| 002-0.6/003-1.5:90/90 | 1.2909575 | 0.6405087 | 0.4542600 | 0.6496417 |
| 003-0.6/003-1.5:90/90 | 1.1496704 | 0.5072949 | 0.2950363 | 0.7679661 |

001 = mdc-STM-001
002 = mdc-STM-002
003 = mdc-STM-003

**Supplementary table S12.** Between formulations comparisons of 10-day LC50 and LC90 values for VK5 mosquitoes fed on treated cattle.

| Comparison | Estimate | Std. Error | t-value | p-value |
| --- | --- | --- | --- | --- |
| 001-0.6/002-0.6:50/50 | 0.9513449 | 0.4334657 | -0.1122468 | 0.9106277 |
| 001-0.6/003-0.6:50/50 | 0.9647061 | 0.4190361 | -0.0842264 | 0.9328765 |
| 001-0.6/003-1.5:50/50 | 0.9823391 | 0.4071531 | -0.0433767 | 0.9654013 |
| 002-0.6/003-0.6:50/50 | 1.0140446 | 0.4935095 | 0.0284586 | 0.9772964 |
| 002-0.6/003-1.5:50/50 | 1.0325793 | 0.4842848 | 0.0672731 | 0.9463643 |
| 003-0.6/003-1.5:50/50 | 1.0182780 | 0.4565668 | 0.0400337 | 0.9680663 |
| 001-0.6/002-0.6:90/90 | 0.9165698 | 0.5121319 | -0.1629076 | 0.8705911 |
| 001-0.6/003-0.6:90/90 | 1.0141546 | 0.4742174 | 0.0298484 | 0.9761880 |
| 001-0.6/003-1.5:90/90 | 1.0997614 | 0.5007874 | 0.1992090 | 0.8420993 |
| 002-0.6/003-0.6:90/90 | 1.1064674 | 0.6179795 | 0.1722831 | 0.8632150 |
| 002-0.6/003-1.5:90/90 | 1.1998665 | 0.6578991 | 0.3037950 | 0.7612841 |
| 003-0.6/003-1.5:90/90 | 1.0844119 | 0.4934883 | 0.1710515 | 0.8641833 |

001 = mdc-STM-001
002 = mdc-STM-002
003 = mdc-STM-003

**Supplementary Table S13.** Lethal concentration (LC50) for KIS and VK5 mosquitoes over 4-day follow up period (4-day LC50). Mean concentrations are given with lower and upper confidence intervals (95% CI).

| Cumulative mortality window (days) | Mortality % | KIS mosquitoes | VK5 mosquitoes | *P*-value (between colony comparison) |
| --- | --- | --- | --- | --- |
| 4 | 50 | 2.89 [2.27-3.67] | 5.26 [4.36-6.35] | 1.0e-07 |
| 4 | 90 | 9.2 [6.72-12.59] | 17.65 [12.86-24.22] | 0.0e+00 |

## Supplementary figures

**Supplementary Fig. S1:**
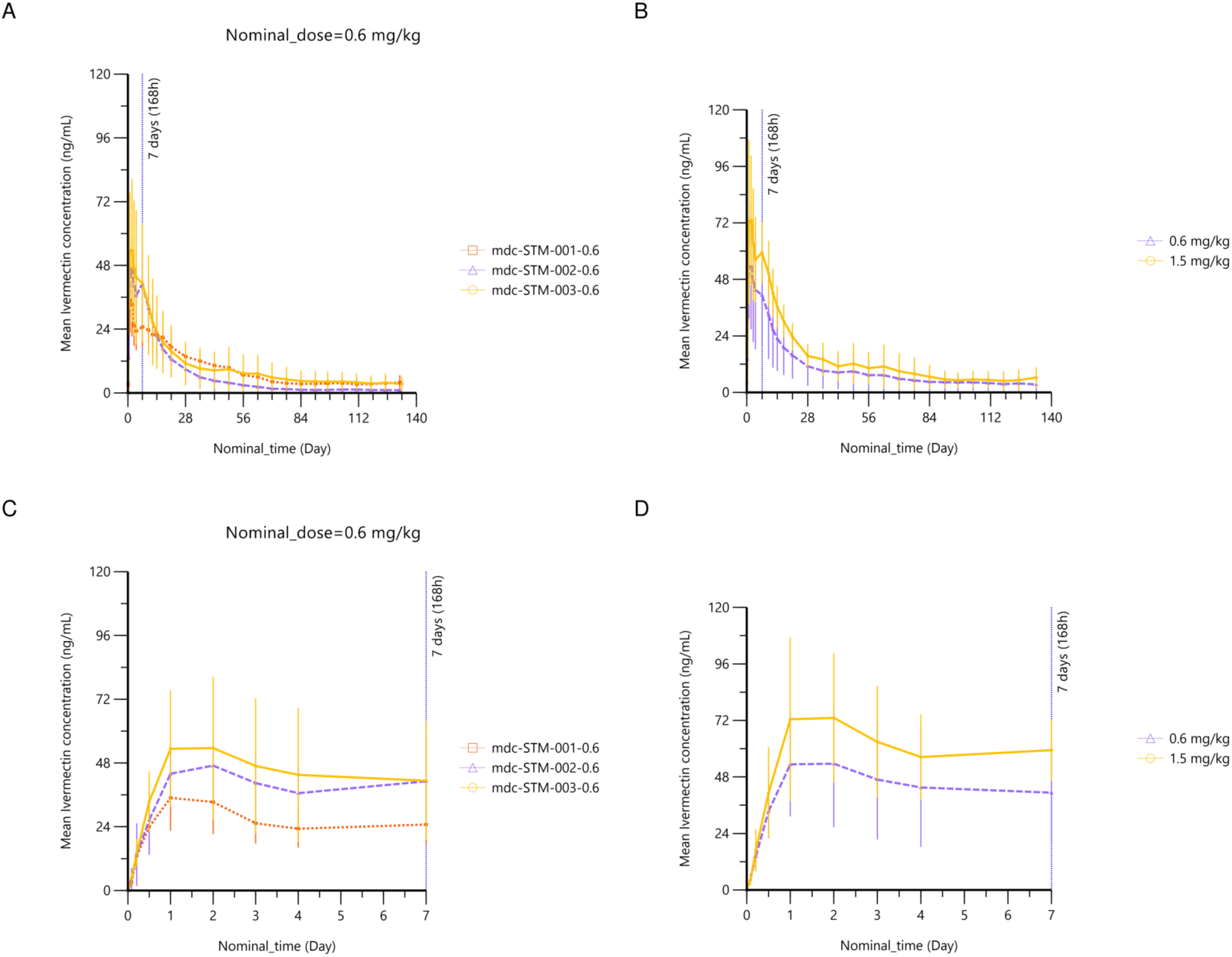
Additional mean concentration–time profiles of ivermectin in cattle plasma. Panels A and C: profiles by formulation at a dose of 0.6 mg/kg for the entire experiment (A) and during the first 7 days post-injection to visualize the initial burst (C). Panels B and D: profiles by dose for the entire experiment (B) and during the first 7 days post-injection (D)

**Supplementary Fig. S2.**
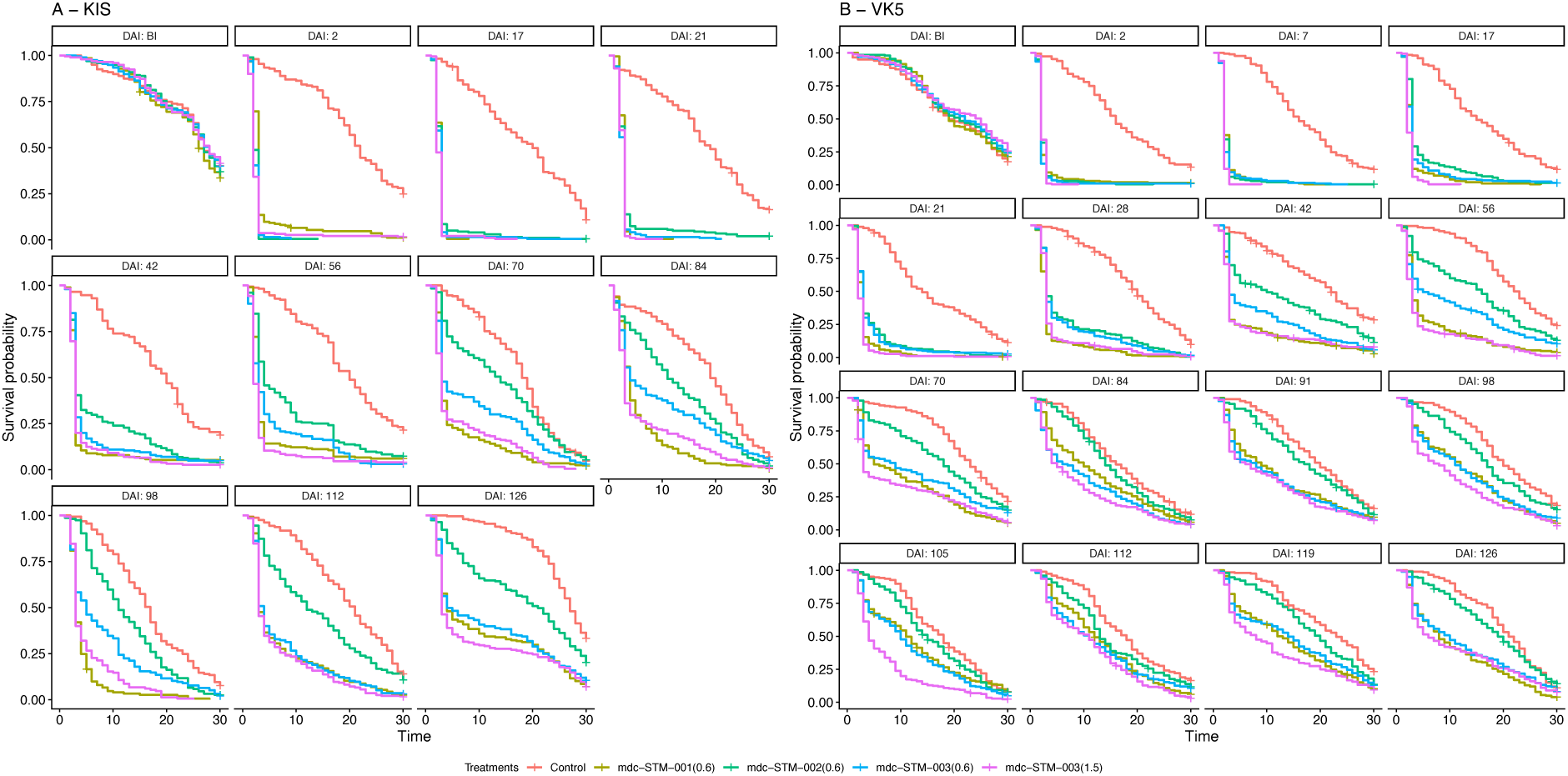
Kaplan-Meier plots illustrating the mosquitocidal effects of the candidate long-lasting ivermectin formulations administered to calves at doses of 0.6 mg/kg (mdc-STM-001, mdc-STM-002 and mdc-STM-003) or 1.5 mg/kg (mdc-STM-003 only). The plots depict the results from each direct skin feeding assay conducted at the indicated time points (Days After Injection, DAI). Mosquito survival was monitored daily for up to 30 days post-blood meal. Pannel A: KIS colony, Pannel B: VK5 colony.

**Supplementary Fig. S3:**
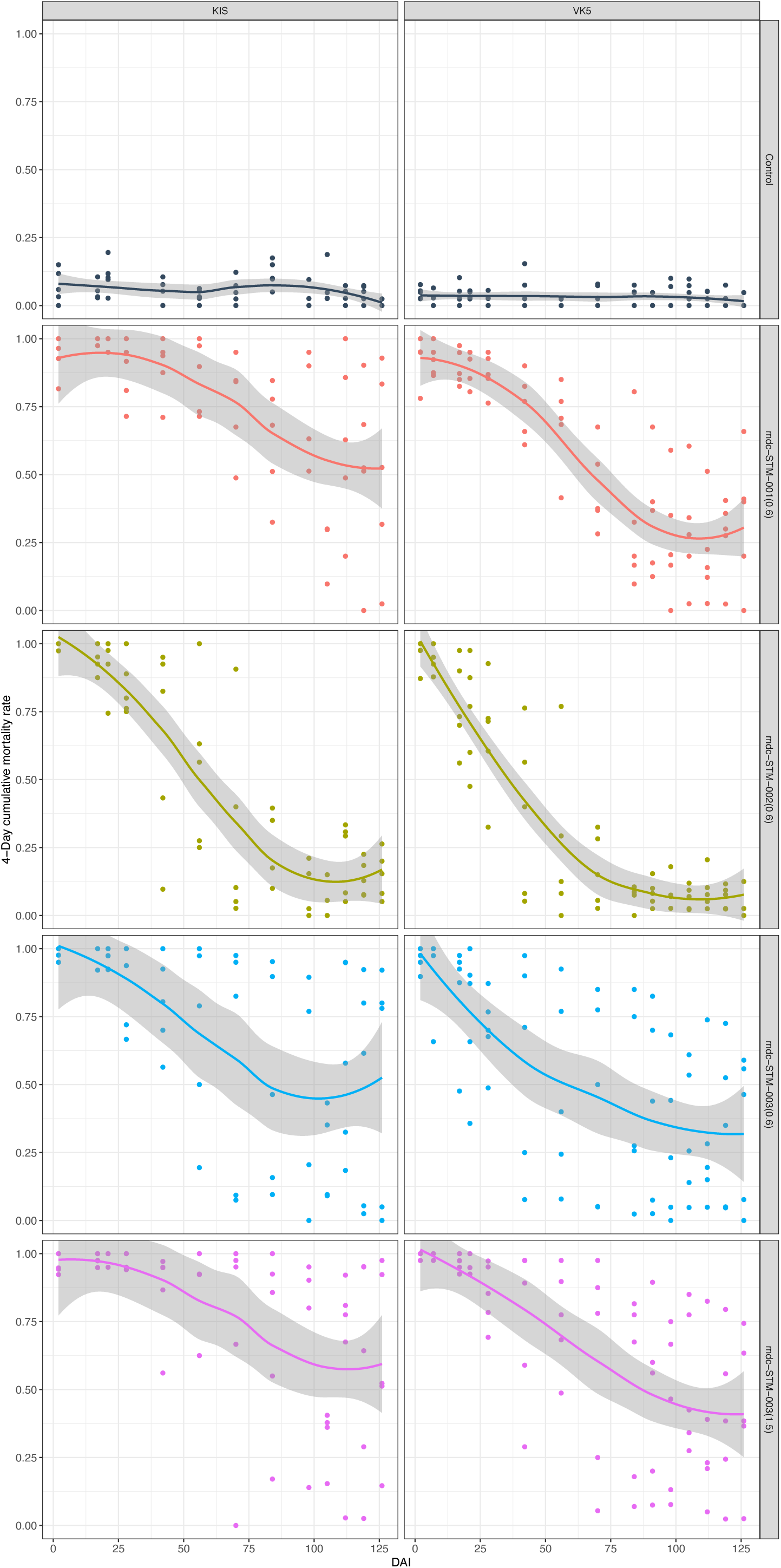
4-day cumulative mortality of KIS and VK5 colony mosquitoes after feeding on calves treated using the candidate formulations. Treated cattle received a single LAIF injection at a dose of 0.6mg/kg for the mdc-STM-001, mdc-STM-002, mdc-STM-003 formulations, and at an additional dose of 1.5 mg/kg for the mdc-STM-003. The smooth line represents mean values estimated by the LOESS (locally estimated scatterplot smoothing) method and grey area shows CI95%. See the main text for further details.

**Supplementary Fig. S4.**
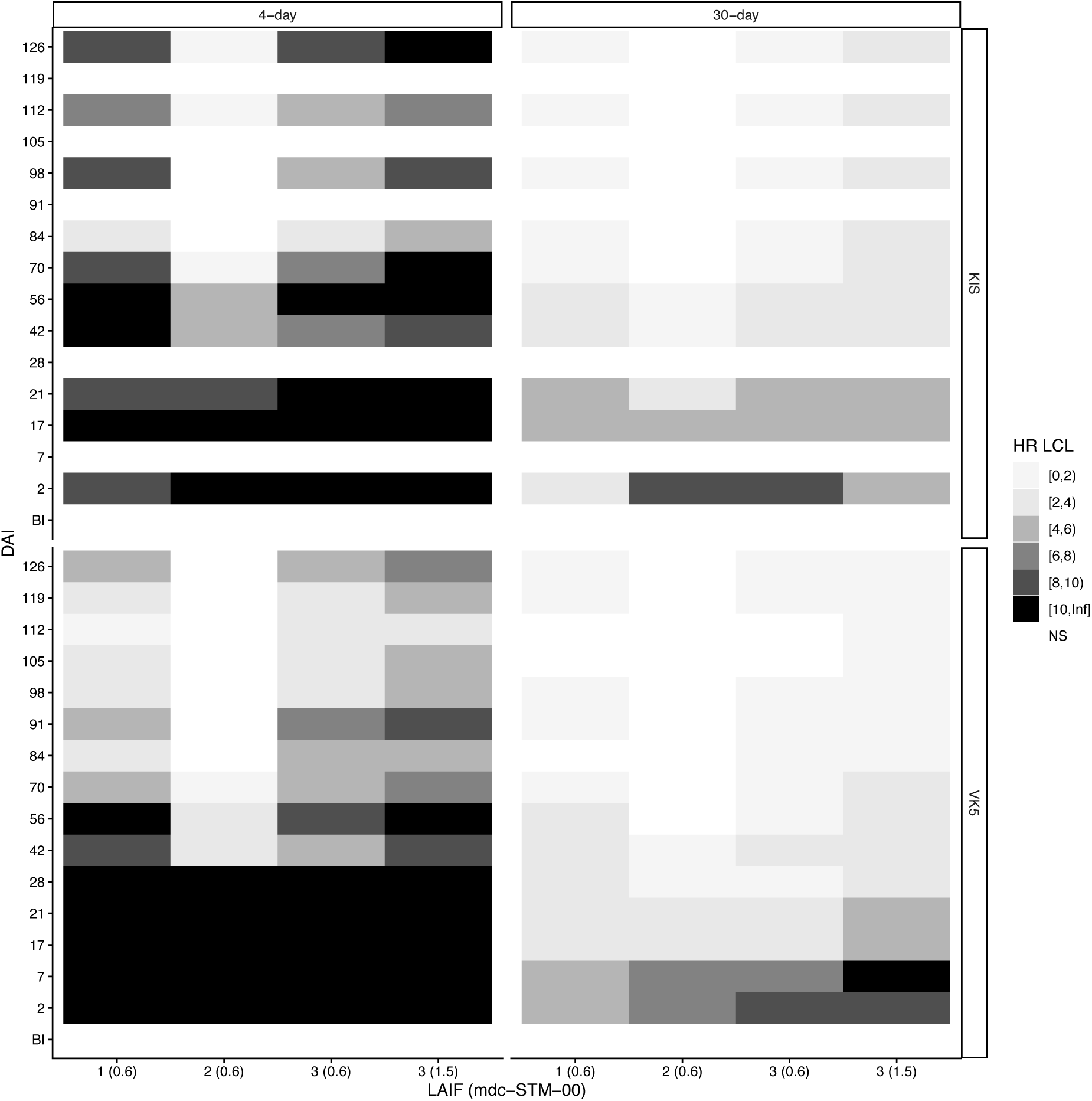
Heat maps illustrating the lower 95% confidence limit (LCL) of 4-day and 30-day mortality hazard ratios (HRs) for KIS and VK5 colony mosquitoes fed at different DAIs on cattle treated with the formulations mdc-STM-001-0.6, mdc-STM-002-0.6, mdc-STM-003-0.6 or mdc-STM-003-1.5. HR and *P-values* for each DAI are available in Additional file 1: Table S7. BI is the timepoint before injection.

**Supplementary Fig. S5.**
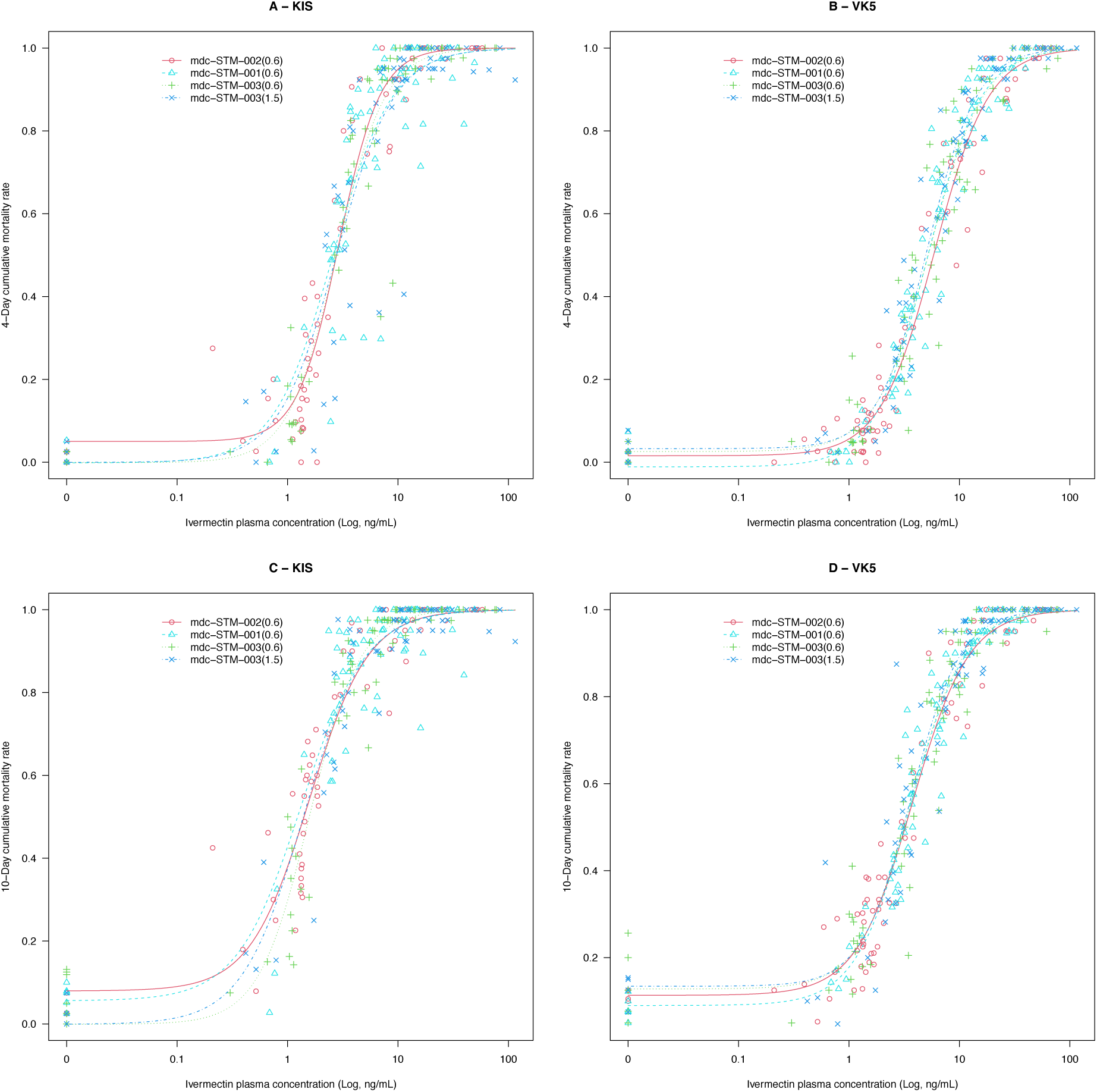
Relationship between ivermectin plasma concentration and 4-days (A and B) or 10-day (C and D) cumulative mortality for each candidate formulation. A and C: KIS colony mosquitoes. B and C: VK5 colony mosquitoes

**Supplementary Fig. 6.**
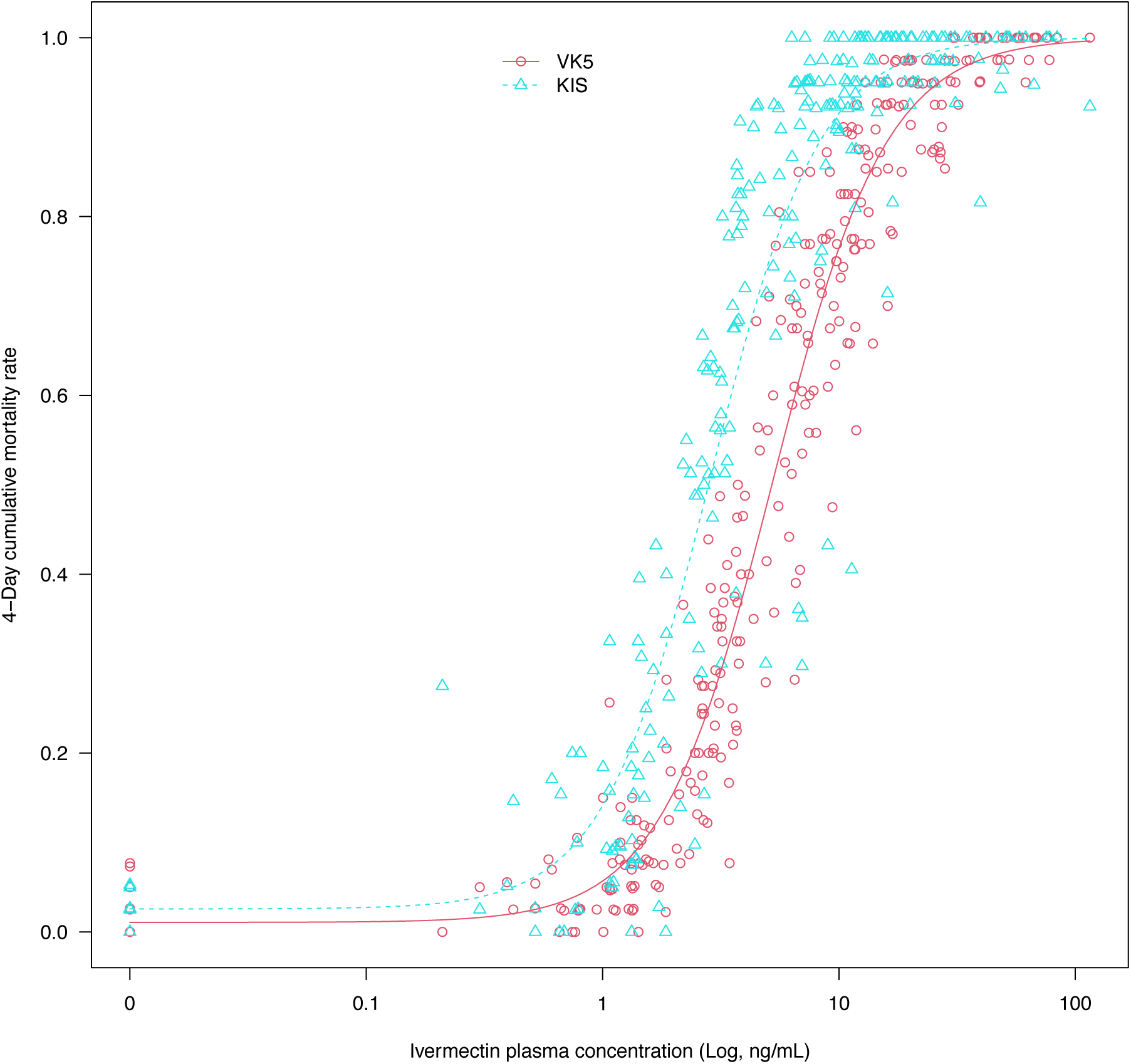
Relationship between ivermectin plasma concentrations and 4-day cumulative mosquito mortality for KIS and VK5 colonies.

